# Creation of a novel zebrafish model with low DHA status to study the role of maternal nutrition during neurodevelopment

**DOI:** 10.1101/2024.07.30.605803

**Authors:** Katherine M. Ranard, Bruce Appel

## Abstract

Docosahexaenoic acid (DHA), a dietary omega-3 fatty acid, is a major building block of brain cell membranes. Offspring rely on maternal DHA transfer to meet their neurodevelopmental needs, but DHA sources are lacking in the American diet. Low DHA status is linked to altered immune responses, white matter defects, impaired vision, and an increased risk of psychiatric disorders during development. However, the underlying cellular mechanisms involved are largely unknown, and advancements in the field have been limited by the existing tools and animal models. Zebrafish are an excellent model for studying neurodevelopmental mechanisms. Embryos undergo rapid external development and are optically transparent, enabling direct observation of individual cells and dynamic cell-cell interactions in a way that is not possible in rodents. Here, we create a novel DHA-deficient zebrafish model by 1) disrupting *elovl2,* a key gene in the DHA biosynthesis pathway, via CRISPR-Cas9 genome editing, and 2) feeding mothers a DHA-deficient diet. We show that low DHA status during development is associated with a small eye morphological phenotype and demonstrate that even the morphologically normal siblings exhibit dysregulated gene pathways related to vision and stress response. Future work using our zebrafish model could reveal the cellular and molecular mechanisms by which low DHA status leads to neurodevelopmental abnormalities and provide insight into maternal nutritional strategies that optimize infant brain health.

## Introduction

Neurodevelopment is dynamic and complex, requiring a coordinated supply of nutrients at precise times and locations. Among these nutrients are lipids, which constitute 50% of the brain’s dry weight, a higher proportion than any other organ (1). Docosahexaenoic acid (DHA, 22:6n-3) is the most abundant long-chain omega-3 polyunsaturated fatty acid (PUFA) in the brain and retina (2–4). DHA primarily resides in the phospholipids of cell membranes and thus helps “shape” each cell’s physical characteristics. However, the importance of DHA in the developing central nervous system (CNS) extends beyond a purely structural role. DHA is crucial for cell signaling, neurotransmission, and neurogenesis, and a lack of DHA may disrupt circuit function and higher order cognitive processes (2,5). Indeed, low DHA status in infancy is associated with an increased incidence of cognitive and psychiatric disorders that develop during childhood, including schizophrenia and autism spectrum disorder (6).

DHA can either be consumed directly (e.g., from marine fish or supplements) or endogenously synthesized through the conversion of the dietary omega-3 PUFA precursor alpha-linolenic acid (ALA, 18:3n-3) (2). Humans have a low DHA biosynthesis capacity (∼5% ALA-to-DHA conversion rate), so offspring rely on maternal DHA transfer, and by extension the maternal diet, to meet their developmental needs (7). Importantly, much of the world now consumes a Western-style diet that is low in DHA and other omega-3 PUFA-rich foods. This dietary shift has also exacerbated the intake imbalance between omega-3 and omega-6 PUFAs. Like omega-3s, omega-6 PUFAs such as arachidonic acid (ARA, 20:4n-6) are essential components of phospholipid cell membranes in the brain (8). However, omega-3s and omega-6s elicit opposing immune-modulatory effects, and an imbalance of omega-3 to omega-6 PUFAs may disrupt the brain’s homeostasis and stress response (9). Notably, DHA supplements are commonly recommended to pregnant and lactating mothers to fill the dietary omega-3 PUFA gap, but concerns remain about maternal DHA status and the neurodevelopmental consequences of DHA insufficiency.

A growing body of evidence supports a role for DHA in the functionality of diverse CNS cell types: In the retina, DHA protects photoreceptors from oxidative stress-induced apoptosis (10) and preserves visual function by facilitating proper photoreceptor disc morphology (11). DHA and its derivatives also have potent anti-inflammatory properties and can affect the activities of microglia, the resident immune cells of the CNS (12). DHA-deficient mice display an increased pro-inflammatory cytokine signature and dysregulated microglia phagocytosis (13,14). The effects of DHA on microglia have been mainly investigated in the context of neurological injury and disease (12). However, during development, microglia also help sculpt the brain’s cellular architecture by releasing growth factors and by engulfing excess myelin, neurons, and synapses (15–17). Therefore, disruptions in microglia-mediated processes could also lead to more extensive neurodevelopmental abnormalities.

Cell culture and rodent studies have constituted the bulk of the evidence thus far, providing valuable insight into DHA functions. However, the cellular and molecular mechanisms of DHA during development, and the role of maternal DHA during this critical period, remain largely unknown. Advancements in the field have been limited by the existing tools and model systems.

Zebrafish are an excellent model for studying neurodevelopmental mechanisms. Embryos undergo rapid external development and are optically transparent, enabling direct observation of individual cells and dynamic cell-cell interactions in a way that is not possible in rodents or cell culture. To leverage the strengths of this powerful model system, we developed a zebrafish model with low DHA status. We used a two-pronged approach, targeting both possible avenues for DHA acquisition: DHA biosynthesis and the maternal diet. To disrupt DHA biosynthesis, we focused on elongation of very long-chain fatty acids protein 2 (*elovl2*), a rate-limiting gene in the DHA biosynthesis pathway. In support of *elovl2* as an ideal target, *Elovl2^-/-^* mice (18,19) and *elovl2* mutant zebrafish (20,21) have reduced systemic DHA levels. We therefore generated a stable *elovl2* mutant zebrafish line using CRISPR/Cas9 technology. We also fed wild-type and heterozygous *elovl2* mothers a DHA-deficient diet, which contained the omega-3 precursor ALA but no preformed DHA.

We found that altering the maternal diet alone decreased wild-type offspring DHA levels, and that offspring DHA status was further reduced by combining maternal genetic and dietary strategies. Interestingly, *elovl2^-/-^*and *elovl2^+/+^* clutchmate siblings had similar DHA levels, suggesting that offspring DHA status largely depends on maternal factors. A proportion of offspring generated from mothers fed the DHA-deficient diet displayed a morphological phenotype, and small malformed eyes were among the abnormal features we observed. Interestingly, vision and stress response gene pathways were still significantly dysregulated in their morphologically normal siblings. These findings align with the known roles of DHA in the retina and strengthen the relevance of our model.

We created a novel zebrafish model with low DHA status by disrupting DHA biosynthesis and modifying the maternal diet. This tool will facilitate novel investigations into how low DHA status alters the cellular and molecular mechanisms guiding neurodevelopment. Addressing this fundamental gap in knowledge may also provide insight into maternal nutritional strategies that optimize infant brain health.

## Materials and methods

### Zebrafish husbandry

All animal work was approved by the Institutional Animal Care and Use Committee at the University of Colorado Anschutz Medical Campus (protocol #370). Adult fish were maintained on a standard 14 h /10 h light/dark cycle. Wild-type zebrafish (*Danio rerio)* were of the AB strain. All offspring used for experiments were generated from pairwise crosses. Embryos were raised in egg water (0.3 g Instant Ocean^®^ sea salt per L ddH_2_O) in 10 cm Petri dishes at 28.5°C, and staged according to hours or days post-fertilization (hpf/dpf) and morphological criteria (22). Prior to experiments, larvae were euthanized via tricaine methanesulphonate (MS-222) in egg water. Sex is undetermined at embryonic and larval stages and was therefore not considered for these experiments.

### Diets

Zebrafish were fed either a commercial lab diet (Gemma Micro; Skretting) containing sufficient levels of DHA (SUFF) or an experimental nutrient-defined DHA-deficient diet (DEF) (**Table 1**). SUFF and DEF differ in their lipid sources: SUFF contains fish meal and fish oil (**Supplemental Table S1**), whereas DEF contains soybean oil (23) (**Supplemental Table S2**). Note that DEF contains only fatty acids with 18 or fewer carbons and 2 or 3 double bonds, which includes ALA, a DHA precursor. The nutrient-defined DEF diet was prepared in 100 g batches at the Oregon State University Sinnhuber Aquatic Research Laboratory as previously described (24) and stored at -20°C until fed to fish. Diet batches were used within 6 months of preparation. When raising zebrafish to adulthood, larvae (∼5 dpf) were added into a rotifer polyculture at ∼275 rotifers/mL, and rotifers were fed with RGComplete™ (Reed Mariculture). A few mL of rotifers and RGComplete™, and a small quantity of SUFF, were added daily. From ∼10 to 21 dpf, juveniles were fed 2-4 times daily with rotifers and SUFF, and were then fed only SUFF from ∼21 to 60 dpf, at which time fish were assigned to either remain on SUFF or transition to the DEF diet. This 60 dpf transition timepoint was based on previous studies (24–26) and the pilot feeding trial described here. All adult fish were fed 2 times/day, with feedings separated by at least 1 h. The adults used to generate offspring for the described experiments had been fed DEF for at least 20 weeks (up to 42 weeks). Because the yolk serves as a zebrafish embryo’s nutrient source until ∼5 dpf, the offspring used for experiments were never fed. The zebrafish model therefore offers a unique closed system for investigating how maternal diet influences offspring development and nutrient status.

**Table 1.** Maternal diet fatty acid composition (g/kg diet)*^a^*.

| **fatty acid** | **SUFF** | **DEF** |
| --- | --- | --- |
| 14:0 | 1.18 ± 0.08 | 0.04 ± 0.00 |
| 16:0 | 13.28 ± 0.55 | 4.72 ± 0.02 |
| 18:0 | 1.30 ± 0.03 | 0.58 ± 0.00 |
| 16:1 | 4.35 ± 0.08 | 0.02 ± 0.00 |
| 18:1 | 1.04 ± 0.04 | 1.30 ± 0.07 |
| 18:2n-6 (LA) | 2.37 ± 0.23 | 4.87 ± 0.28 |
| 18:3n-3 (ALA) | 1.21 ± 0.10 | 2.38 ± 0.08 |
| 20:4n-6 (ARA) | 0.81 ± 0.00 | 0.01 ± 0.00 |
| 20:5n-3 (EPA) | 6.63 ± 0.54 | 0 |
| 22:6n-3 (DHA) | 7.61 ± 0.30 | 0 |
| LA:ALA | 1.96 | 2.05 |
| ARA:DHA | 0.11 | 6.67 |

<sup>a</sup>fatty acids were analyzed by GC/MS for representative diet samples (n=3 replicates/diet). Values reported as mean ± SEM. ALA, alpha-linolenic acid; ARA, arachidonic acid; DEF, DHA-deficient diet; DHA, docosahexaenoic acid; EPA, eicosapentaenoic acid; LA, linoleic acid; SUFF, DHA-sufficient diet.

### Pilot feeding trial growth assessments

A pilot feeding trial was conducted to ensure that wild-type fish fed the experimental DEF diet (WT-DEF) grew similarly to fish fed the SUFF diet (WT-SUFF). This feeding trial was also used to determine what age to transition fish onto the DEF diet during development (30 or 60 dpf) for the wild-type and *elovl2* mutant study cohorts. Fish were fed ∼2.5% body mass/day, divided equally across 4 feedings/day for fish that were 1-2 months of age and 2 feedings/day for fish that were ≥2 months old. The total tank wet mass at the beginning of the pilot study was used for initial diet calculations, and diet quantities were adjusted every 2 weeks as needed to account for fish growth. The diet quantities needed for each tank were portioned into microcentrifuge tubes and stored at -20°C until fed to fish. The primary outcomes for the pilot feeding trial were fish body mass and length, which were recorded every 2 weeks. Average individual fish body mass was calculated by dividing the measured total tank wet mass by the number of fish per tank. Fish lengths (nose to caudal peduncle) were traced from aerial tank photos using an ImageJ plugin, and measurements were calibrated to a ruler placed in the tank.

### Generation of elovl2 mutant lines

*elovl2* mutant lines were generated using CRISPR/Cas9-mediated genome editing as described previously (27), with slight modifications. Briefly, using the IDT Alt-R System, CRISPR/Cas9 RNP complexes were prepared using two predesigned IDT Alt-R CRISPR-Cas9 crRNAs (Dr.Cas9.ELOVL2.1.AA and Dr.Cas9.ELOVL2.1.AB), Alt-R tracrRNA, and Alt-R S.p. Cas9 nuclease V3. A dual-injection strategy was employed using the selected crRNAs. The lyophilized crRNAs were first reconstituted in IDTE buffer (100 µM) and mixed with tracrRNA (100 µM) and Nuclease-Free Duplex Buffer (IDT) to prepare a 3 µM gRNA solution. The solution was then heated to 95°C for 5 min, cooled to room temperature, and stored at -20°C until the day of injections. For RNP complex assembly, the gRNA solution was combined with an equal volume of 3 µM Cas9 diluted in Cas9 working buffer (20 mM HEPES, 150 mM KCl, pH 7.5), incubated at 37°C for 10 min, and then microinjected into zebrafish embryos at the one-cell stage (approximately 3 nL per embryo).

Mutations causing insertions and deletions were confirmed in 1 dpf injected embryos via PCR amplification and gel electrophoresis (>60% cutting efficiency) using primers that flanked the gRNA target sites, herein described as target A (within exon 3) and target B (within exon 4). Primer sequences are listed in **Supplemental Table S3**. Stable, germline transformed, *elovl2* mutant lines were established by raising injected embryos to adulthood and outcrossing F0 founders (confirmed via fin biopsy, PCR amplification, and gel electrophoresis) with wild-type zebrafish. To verify germline transmission, F1 progeny from each clutch were screened at 1 dpf using the target A and target B primers. To identify mutant alleles, Sanger sequencing was performed on F1 embryos that displayed indels at each target as assessed by gel band patterns. Prior to sequencing, the amplified PCR was purified using the QIAquick PCR Purification Kit (QIAGEN) according to the manufacturer’s instructions. F1 offspring with mutations predicted to prematurely terminate protein translation were selected for raising and establishing the stable mutant lines. The two selected mutant alleles were: 157 bp deletion at target A and 13 bp insertion at target B (*elovl2^co100^*) (**Figure 4**; **Supplemental Figure S2**); 4 bp and 10 bp deletions at targets A and B, respectively (*elovl2^co101^*) (**Supplemental Figures S1 and S2**). Unless otherwise noted, the *elovl2* mutant offspring used in these experiments were generated from pairwise matings of heterozygous *elovl2^co100^*fish fed DEF.

### elovl2 genotyping

DNA lysates were prepared for larvae and adults by digesting a small piece of tail/fin tissue in NaOH (50 mM), incubating at 95°C for 10 min, adding 1M Tris-HCl (pH 8.0) at a 10:1 NaOH to Tris-HCl ratio, vortexing briefly, and then storing at 4°C. For genotyping, GoTaq^®^ Green Master Mix (Promega) was combined with the appropriate forward and reverse primers (10 µM), nuclease-free water, and DNA lysate. PCR amplification was performed using a T100 Thermal Cycler (Bio-Rad) at 95°C for 3 min, 34 cycles of 95°C for 30 s, 52°C for 30 s, and 72°C for 30 s, followed by 72°C for 5 min. Upon establishing the *elovl2^co100^* and *elovl2^co101^* mutant lines, allele-and target-specific genotyping primers were designed to allow for better resolution of mutant and wild-type bands on a 1.5% agarose / 1.5% metaphor gel. Oligonucleotide sequences and the wild-type and mutant allele amplicon sizes are listed in **Supplemental Table S3**, and example gels are shown in **Supplemental Figure S2**. All genotyping primer sets were designed using Primer3 (https://primer3.ut.ee).

### RNA fluorescent in situ hybridization

RNA-fluorescent *in situ* RNA hybridization (RNA-FISH) was performed on 4 dpf WT-SUFF larvae using the RNAScope Multiplex Fluorescent V2 Assay Kit (ACD). Larvae were fixed in 4% paraformaldehyde in 1X DEPC PBS, rocking O/N at 4°C. Larvae were then embedded in 1.5% agar/5% sucrose blocks and immersed in 30% sucrose O/N at 4°C. Blocks were frozen on dry ice and sectioned on a cryostat microtome (15µm thick transverse sections, collected on polarized microscope slides). FISH was performed using the manufacturer’s protocols with minor modifications. Slides were covered with Parafilm for all incubation steps to maintain moisture and disperse reagents across all tissue sections. Probes for zebrafish *elovl2*-C1 and *slc1a3b*-C2 were designed and synthesized by the manufacturer and used at 1:50 dilutions. Transcripts were fluorescently labeled with Opal 520 (1:1500) and Opal 570 (1:500) using the Opal 7 Kit (Perkin Elmer; NEL797001KT). Images were acquired on a Zeiss LSM 880 inverted confocal system with 20X air and 40X oil objectives using Zen Black software (Carl Zeiss). Fiji/ImageJ was used to adjust brightness and contrast, generate standard deviation z-projections, and to split or merge channels. RNA-FISH figures were created using Adobe InDesign, with Adobe Photoshop color balance adjustments (Levels tool, “auto” setting).

### Lipidomics

Lipid analyses were conducted in zebrafish offspring generated by WT-SUFF, WT-DEF, and *elovl2*-DEF mothers, as well as representative samples of the SUFF and DEF diets. Offspring were collected immediately postspawning (0 hpf) or as larvae (4 dpf) into screw-cap microcentrifuge tubes, flash frozen in liquid nitrogen, and stored at -80°C. For genotyping purposes, tail-clips were performed on all 4 dpf larvae prior to collections. Note that although offspring from WT clutches were not genotyped, tail-clips were still performed on these fish to standardize the amount of tissue collected.

### Total fatty acid analysis (GC/MS)

For total fatty acid analyses, individual offspring and diet samples were homogenized in 200 µL Optima™ water for 2 x 2 min at 25 Hz using a TissueLyser bead mill (QIAGEN), and then an aliquot of methanol (200 µL) was added. Diet and zebrafish offspring homogenates were saponified with 0.5N NH_4_OH in 50% MeOH for 2h at 37°C and combined with a mixture of stable isotope-labeled fatty acid standards [d_2_]myristic, [^13^C_4_]palmitic, [d_4_]stearic, [d_8_]arachidonic and [d_5_]docosahexaenoic acids. After saponification, samples were acidified and extracted using isooctane. Samples were then dried and derivatized with pentafluorobenzyl bromide in N,N-diisopropylethylamine as described previously (28). Analysis of the samples was performed by negative ion chemical ionization GC/MS on a Finnigan DSQ GC/MS system (Thermo Finnigan, Thousand Oaks, CA). The mass spectrometer was operated in the negative ion chemical ionization mode using methane as reagent gas. Data were acquired by selected ion monitoring of the following fatty acids: lauric (m/z 199), myristic (m/z 227), palmitic (m/z 255), stearic (m/z 283), linolenic (m/z 277), linoleic (m/z 279), oleic (m/z 281), eicosapentaenoic (m/z 301), arachidonic (m/z 303), and docosahexaenoic acid (m/z 327). The ions at m/z 229, 259, 287, 311 and 332 were monitored for [d2]myristic, [^13^C_4_]palmitic, [d_3_]stearic, [d_8_]arachidonic and [d5]docosahexaenoic acids, respectively. Concentration was determined using stable isotope dilution with standard curves generated for each free fatty acid. Results are reported as total µg fatty acid per embryo.

### Phospholipid analysis (LC/MS/MS)

Phospholipid (PL) analysis was conducted on a subset of pooled zebrafish offspring to assess the distribution of fatty acids across PL classes and species of interest. Samples were homogenized in 300 µL Optima™ water for 2 x 2 min at 25 Hz using a TissueLyser bead mill (QIAGEN). Another 650 µL Optima™ water was added, and 750 µL of this total homogenized sample was used as the starting sample volume for lipid extractions. Methanol (900 µL) and an internal standard cocktail containing PC-19:0/19:0 (2000 pmol), d7-PC-18:1/OH (200 pmol), and 1,2,3-triheptadecanoyl glycerol (1500 pmol) was added. The lipid extraction was then performed with the addition of methyl-tert-butyl ether (3 mL) according to Matyash et al (29). Samples were injected into an HPLC system connected to a triple quadrupole mass spectrometer (Sciex 3200, Framingham, MA) and normal phase chromatography was performed using a silica column (150x2 mm, Luna Silica 5 µm, Phenomenex). The mobile phase system consisted of solvent A (isopropanol/hexane/water (58/40/2, v/v)) and 35% solvent B (hexane/isopropanol/water (300/400/84, v/v/v) both containing 10 mM ammonium acetate. Mass spectrometric analysis was performed in the negative ion mode using multiple reaction monitoring (MRM). The precursor ions monitored were the molecular ions [M-H]^-^ for PS, LPS, PE, LPE, PI, and LPI and the acetate adducts [M+CH_3_COO]^−^ for PC and LPC. The product ions analyzed after collision-induced decomposition were the carboxylate anions corresponding to the acyl chains. Quantitative results were determined using stable isotope dilution with standard curves for saturated and unsaturated phospholipid compounds.

The PL analysis included only a subset of larval samples (n=3 pooled samples/group, n=3 larvae/pool). Due to the low sample numbers, all PLs containing the same fatty acid moiety were grouped together (i.e., DHA-PLs and ARA-PLs) for statistical analysis. The following comparisons were then analyzed for each PL grouping using Mann-Whitney tests: WT-SUFF versus WT-DEF; WT-SUFF versus *elovl2*-DEF; and *elovl2^+/+^* versus *elovl2^-/-^.* The *elovl2-* DEF group included both the *elovl2*^+/+^ and *elovl2*^-/-^ offspring samples.

### RNA isolation and bulk RNA-sequencing

RNA was isolated from pooled 4 dpf WT-SUFF and *elovl2*-DEF larvae using the RNeasy^®^ Plus Micro Kit (QIAGEN) according to the manufacturer’s instructions. For homogenization, samples were passed 20 times through a 23-gauge needle attached to a 1 mL syringe. Each pooled sample contained offspring from a single clutch, representing an individual mother (n=25 larvae/pool, n=3 replicates/group). RNA purity, quantity, and integrity were determined with NanoDrop™ (ThermoFisher Scientific) and TapeStation 4200 (Agilent) analysis. All samples passed the quality control tests prior to RNA-seq library preparation. Sample libraries were constructed using the Universal Plus™ mRNA-Seq library preparation kit with NuQuant^®^ (Tecan), using an input of 200 ng total RNA. Libraries were sequenced on a NovaSeq X (Illumina) at ∼40M read pairs/80M total reads per sample with a 2x150bp configuration. Raw sequencing reads were de-multiplexed using bcl2fastq.

### RNA-sequencing analyses

For our bulk RNA-seq, derived sequences were analyzed with the nf-core rnaseq pipeline v.3.12.0 (30,31) mapping reads to the GRCz11 zebrafish genome using STAR and Salmon. The resulting raw count matrix was analyzed with the nf-core differentialabundance pipeline v.1.4.0 (30,32) using filtering_min_abundance = 10 and filtering_min_samples = 3 to generate a shinyngs app for the contrast between the 3 WT-SUFF and 3 *elovl2*-DEF samples. A volcano plot showing the differentially expressed genes was generated with Pluto (https://pluto.bio). The false discovery rate (FDR) method was applied for a multiple testing correction. Significance thresholds were set using an FDR-adjusted *P-*value (q<0.1) and ≥2 fold-change. Gene ontology analysis was conducted with Metascape (www.metascape.org) (33) using the list of significantly differentially expressed genes (110 up; 180 down). *P*-values were calculated based on the cumulative hypergeometric distribution, and Benjamini-Hochberg procedure was used to account for multiple comparisons.

Published single-cell RNA-seq datasets from zebrafish, mouse, and human tissues were evaluated for *elovl2/Elovl2/ELOV2* expression, respectively. Mouse and human expression data were sourced from the CZ CELLxGENE Discover platform (https://cellxgene.cziscience.com/gene-expression), which provides aggregated transcriptomic data from a collection of community-contributed studies (34). To assess *Elovl2*/*ELOVL2* expression across all tissues and determine relative expression levels in the liver, brain, and eye specifically, all available datasets on the platform (42 mouse and 186 human) were used in an organism-specific analysis. The platform’s datasets were then also filtered by tissue-type to assess relative expression levels across cell-types in a particular tissue. Mouse *Elovl2* expression results were curated from 2 liver, 1 eye, and 15 brain datasets, and the human *ELOVL2* expression results included 23 liver, 25 eye, and 99+ brain datasets. The results and dot plots for these analyses were downloaded from the CZ CellxGene Discover platform, and the dot plots were incorporated into a summary figure for this manuscript using Adobe Illustrator. Zebrafish *elovl2* expression data for 4 dpf larvae were acquired from the Daniocell online resource (https://daniocell.nichd.nih.gov/index.html) and associated datasets (35,36). As with the mouse/human data, zebrafish expression was assessed across all tissues first, followed by cell-type-specific evaluations for the liver, brain, and eye. Daniocell separates cell-types into multiple clusters, so a single representative cluster for each cell-type. These clusters were as follows: endo.1, endo.6, neur.24, glia.19, microglia, eye.1, eye.28, glia.24. A cluster identified as radial glial cells (glia.27) was used in lieu of an astrocyte cluster, as radial glia give rise to astrocytes during development. The relevant dot plots were then exported from the Daniocell online resource. To better visualize the zebrafish, mouse, and human expression patterns side-by-side, Adobe Illustrator was used to match the expression color gradients and dot size scales between the zebrafish and mouse/human dot plots.

### Larval morphological assessment

Visible morphological phenotypes were assessed in clutches generated for the lipidomics and RNA-seq experiments. At 4 dpf, four abnormal features were used to characterize morphological phenotypes. These features included: small or malformed eyes, curved body axis, pericardial edema, and uninflated swim bladder. If larvae exhibited ≥3 of these features, they were binned into the “severe” phenotype category. Representative larvae were imaged using a Leica stereoscope and SwiftImaging software for Mac (version 3).

### Quantification and statistical analysis

Graphs, figures, and other visualizations were generated using GraphPad Prism for Mac (version 10) and Adobe Software (InDesign; Illustrator; Photoshop). Except for the bulk RNA-seq analysis, statistics were performed in GraphPad Prism. Normality and homogeneity of variance were evaluated using the Shapiro-Wilk and Brown-Forsythe or F tests, respectively. When data met these assumptions, Student’s *t-*tests, 1-way ANOVAs, or 2-way ANOVAs were used to test for differences in means between experimental groups. When data did not exhibit normal distributions or homogeneity of variance, the appropriate non-parametric tests were performed.

## Results

### Wild-type zebrafish fed a DHA-deficient diet have similar growth trajectories as fish fed DHA-sufficient diet

Our main objective was to generate zebrafish offspring with low DHA status by altering the maternal diet and DHA biosynthesis capacity. Toward this goal, we first obtained a DHA-deficient diet (DEF) and verified that DEF-fed juvenile zebrafish grew similarly to siblings fed a commercially available chow that reliably promotes zebrafish growth (37). Although zebrafish nutrient requirements have yet to be defined (38), we deemed the chow to have sufficient DHA because DHA concentrations were comparable to previously published zebrafish study diets (39) (**Table 1**). Thus, we refer to this chow as the DHA-sufficient diet (SUFF) throughout our experiments.

We fed SUFF or DEF to zebrafish starting at either 30 or 60 dpf, and then recorded body mass and length every 2 weeks for 8 weeks (**Figure 1**). As expected, juvenile zebrafish grew significantly during the feeding period, regardless of whether diets were initiated at 30 dpf (body mass, *P*=0.0048; length, *P*<0.001) or 60 dpf (body mass, *P<*0.001; length*, P*<0.001). Growth did not differ between the SUFF-and DEF-fed fish for either cohort (body mass: 30 dpf, *P*=0.454 and 60 dpf*, P*=1.00; length: 30 dpf, *P=*0.191 and 60 dpf*, P=*0.859), and no adverse health effects were evident in either diet group. From this pilot feeding trial, we concluded that DEF is suitable for rearing zebrafish, and we therefore used this diet as part of our strategy to generate offspring with low DHA status. Study parents were transitioned onto DEF at approximately 60 dpf and were fed this diet throughout all experiments. A group of wild-type fish remained on SUFF throughout adulthood, and their offspring served as a reference control group (WT-SUFF).

**Figure 1.**
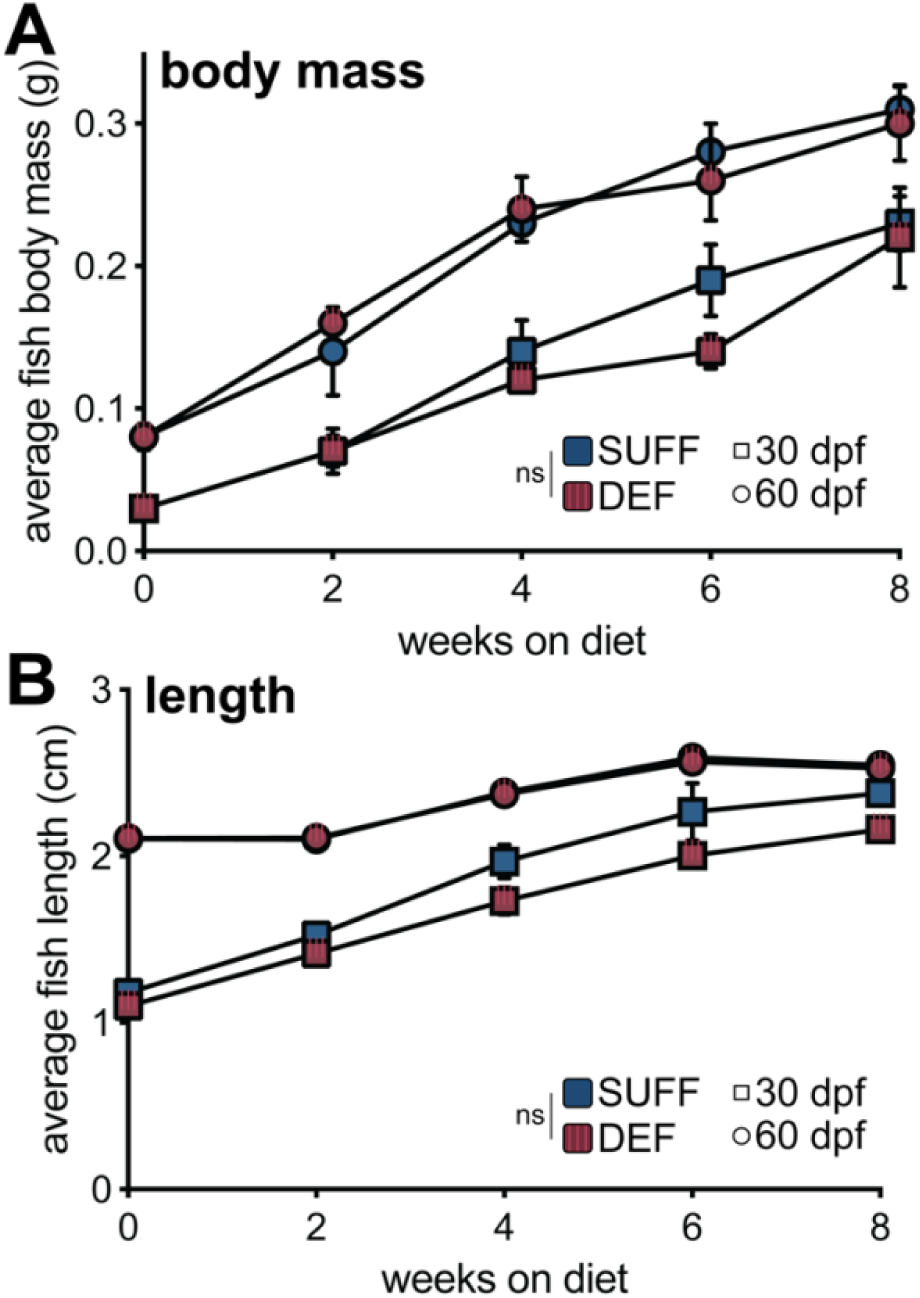
Wild-type zebrafish fed a DHA-deficient diet have similar growth trajectories as fish fed DHA-sufficient diet. **(A)** Average fish body mass, calculated by dividing total tank wet mass by the number of fish per tank. **(B)** Average fish length (nose to caudal peduncle), calibrated to ruler in tank. Separate cohorts of fish were fed the assigned diets starting at either 30 or 60 dpf, indicated in the plots by square or circle symbols, respectively. The main effects of diet and time (weeks on diet) were analyzed by 2-way repeated measures ANOVAs for the 30 and 60 dpf cohorts separately. Body mass and length did not differ between the SUFF and DEF groups for either initiation age. As expected for developing zebrafish, body mass and length significantly increased with feeding duration (body mass: 30 dpf, *P*=0.0048 and 60 dpf*, P<*0.001; length: 30 dpf, *P*<0.001 and 60 dpf*, P*<0.001). Values are mean ± SEM. DEF, DHA-deficient diet; SUFF, DHA-sufficient diet; dpf, days post fertilization.

### Zebrafish central nervous system tissues express elovl2 during development

*elovl2* is a key gene in the DHA biosynthesis pathway (40). In zebrafish, *elovl2* is required for the conversion of eicosapentaenoic acid (EPA, 20:5n-3) to docosapentaenoic acid (DPA, 22:5n-3), which can then be converted to DHA (41). The liver is the primary site for lipid metabolism and DHA biosynthesis, and thus expresses *elovl2* at high levels. However, we wondered whether cells in the developing zebrafish central nervous system (CNS) also express *elovl2*. Local DHA synthesis could guarantee an adequate and accessible DHA supply during neurogenesis and other critical neurodevelopmental processes. Therefore, we first investigated the expression patterns of *elovl2* (zebrafish), *Elovl2* (mouse), and *ELOVL2* (human) in CNS tissues using publicly available single-cell RNA-seq datasets (**Figure 2**). Mouse and human brain astrocytes, and to a lesser degree, oligodendrocytes and neurons, express *Elovl2/ELOVL2* (42–48). Similarly in developing zebrafish, radial glial cell populations, from which astrocytes are derived, express *elovl2* throughout development, starting as early as 14-21 hpf (35,36). In the murine and human eye, retinal cone and rod photoreceptor cells also express *Elovl2/ELOVL2* (49–52), whereas Müller glia cells are one of the top *elovl2-* expressing cell types in the developing zebrafish eye (35,36).

**Figure 2.**
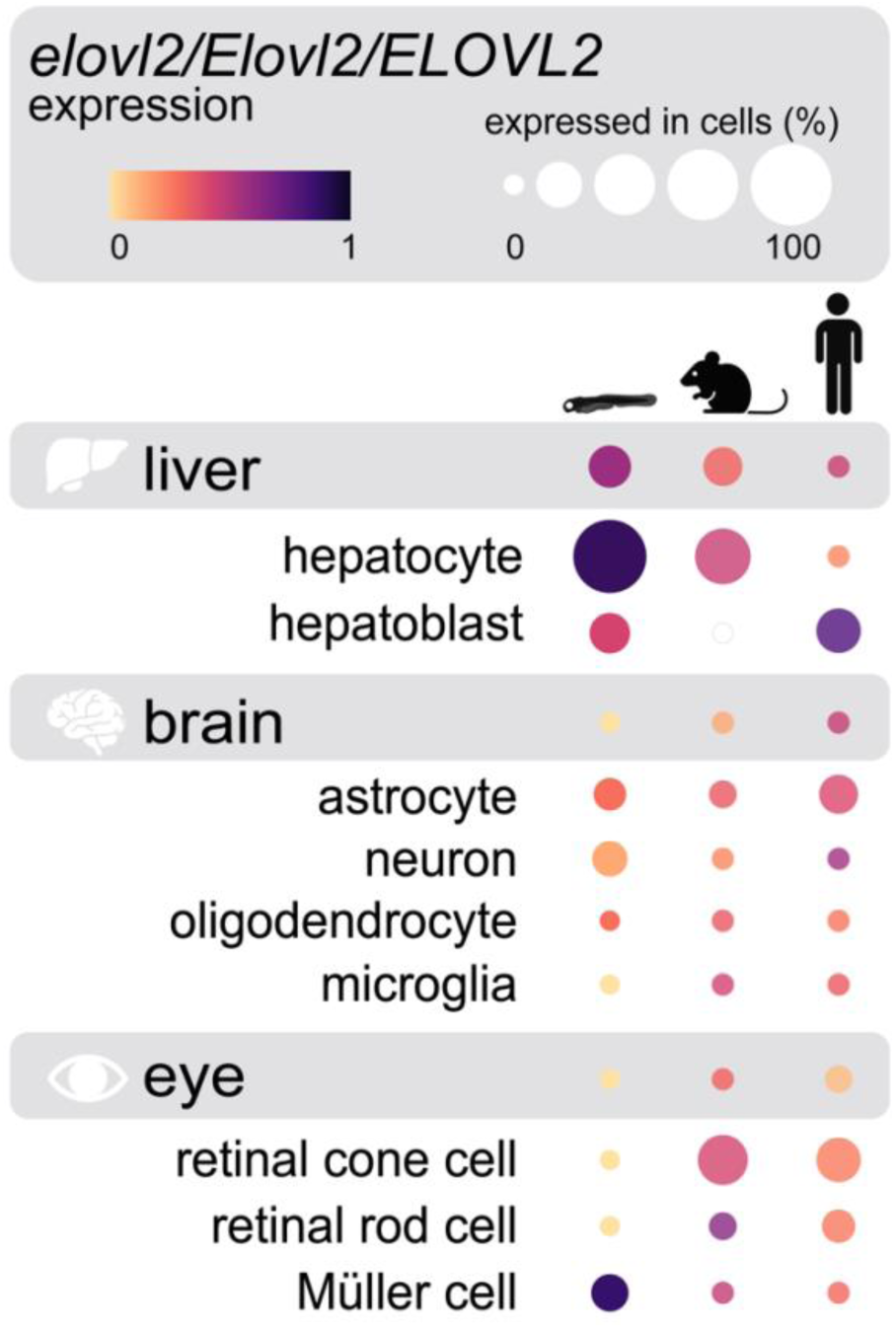
Zebrafish, mouse, and human central nervous system tissues express *elovl2/Elovl2/ELOVL2*, respectively, a critical gene in the DHA biosynthesis pathway. In all three organisms, liver cells strongly express *elovl2/Elovl2/ELOV2*, whereas brain and eye cell-types express this gene at comparatively lower levels. Dot colors represent normalized expression levels (ranging from 0 to 1) for a particular organism, tissue, or cell-type. For each organism, expression comparisons can be made between tissues, e.g., liver versus brain (grey boxes), as well as between cell-types within a given tissue, e.g., astrocyte versus neuron. Dot sizes are scaled to indicate the percentage of each cell-type that expressed the gene of interest. Mouse and human expression patterns are from aggregated datasets submitted to the CZ CELLxGENE Discover platform (doi:10.1101/2023.10.30.563174; https://cellxgene.cziscience.com/gene-expression). Zebrafish expression data (4 dpf larvae) are sourced from the Daniocell datasets and online resource (https://daniocell.nichd.nih.gov/index.html). Schematic design was inspired by the CZ CELLxGENE Discover online portal. Organism and tissue images were created using BioRender.com.

To validate these *elovl2* expression patterns, we performed RNA-FISH in wild-type larvae at 4 dpf, an age when neurodevelopmental processes such as myelination are underway. We detected *elovl2* transcripts in the eye, brain, and spinal cord (**Figure 3**). Specifically in the brain and spinal cord, *elovl2* puncta colocalized with *slc1a3b* transcripts, which mark astrocytes, corroborating the single-cell RNA-seq datasets. Although not quantified, *elovl2* transcripts appeared to be more abundant throughout the brain compared to the spinal cord. Taken together, these data show that *elovl2* is expressed in the developing CNS, indicative of local DHA production.

**Figure 3.**
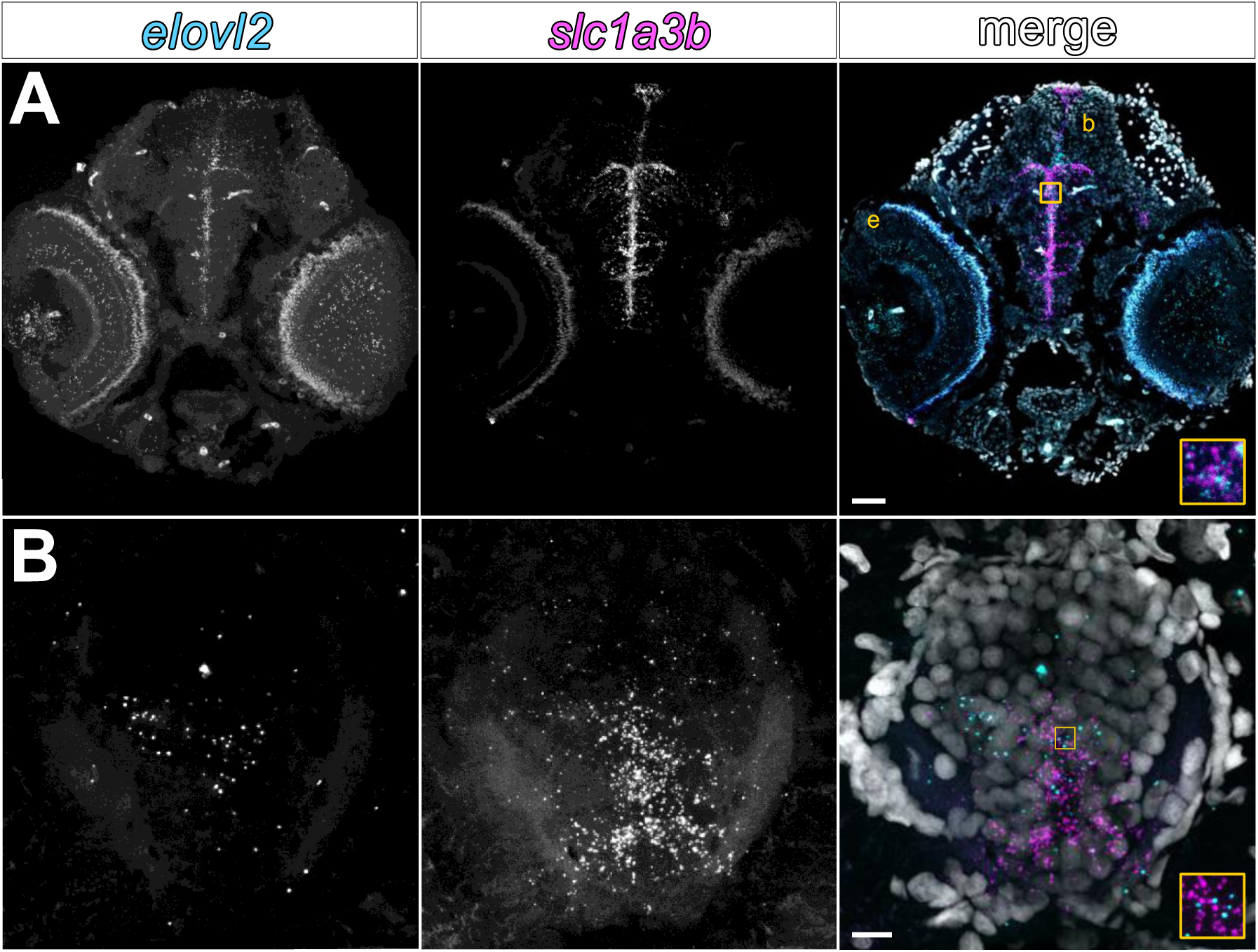
zebrafish central nervous system tissues express *elovl2* during development. RNA Fluorescent *in situ* hybridization performed on transverse sections of 4 dpf wild-type larvae. **(A)** Head, showing *elovl2* puncta in the eye (e) and brain (b), 50 µm scale bar. **(B)** Spinal cord, 10 µm scale bar. Colocalization of *elovl2* and *slc1a3b* (astrocyte/radial glia cell marker) transcripts in the brain and spinal cord supports existing single-cell RNA-seq data.

*Elovl2^-/-^* mice (18,19) and *elovl2* mutant zebrafish (20,21) have decreased systemic DHA levels compared to their wild-type counterparts. During DHA deficiency, *Elovl2^-/-^* mice exhibit an altered immunophenotype of bone-derived macrophages, with an upregulation of pro-inflammatory markers in M1 macrophages and a downregulation of anti-inflammatory markers in M2 macrophages (53). Notably, the number and morphological complexity of the CNS-resident macrophages known as microglia are also increased in *Elovl2^-/-^* mice (13). These findings support *elovl2* gene disruption as a viable strategy to model DHA deficiency and evaluate the neural and immune-related consequences. We therefore generated *elovl2* mutant zebrafish using CRISPR/Cas9 technology. We used a dual-guide RNA approach, targeting exons 3 and 4, which we refer to as targets A and B, respectively (**Figure 4A**). We recovered an allele (*elovl2^co100^*) with a 157 bp deletion at target A and 13 bp insertion at target B (**Figure 4B**). The *elovl2^co100^* allele results in an early stop codon in exon 5, with a predicted truncated protein of 80 AA compared to the 260 AA wild-type Elovl2 (**Figure 4C**). We also recovered a second *elovl2* mutant allele (*elovl2^co101^*) with 4 bp and 10 bp deletions at targets A and B, respectively (**Supplemental Figures S1 and S2**). The *elovl2^co101^*allele results in an early stop codon in exon 3, predicting a premature termination of protein translation (55 AA). We established stable mutant lines for both alleles and determined that they are homozygous viable. We also confirmed that fatty acid profiles were similarly reduced in *elovl2^co100^* and *elovl2^co101^* mutant larvae (**Supplemental Table S4; Supplemental Figure S3**). We opted to use the *elovl2^co100^* mutant line due to ease of genotyping, so for all experiments, *elovl2* mutant offspring were generated from pairwise matings of heterozygous *elovl2^co100^* parents.

**Figure 4.**
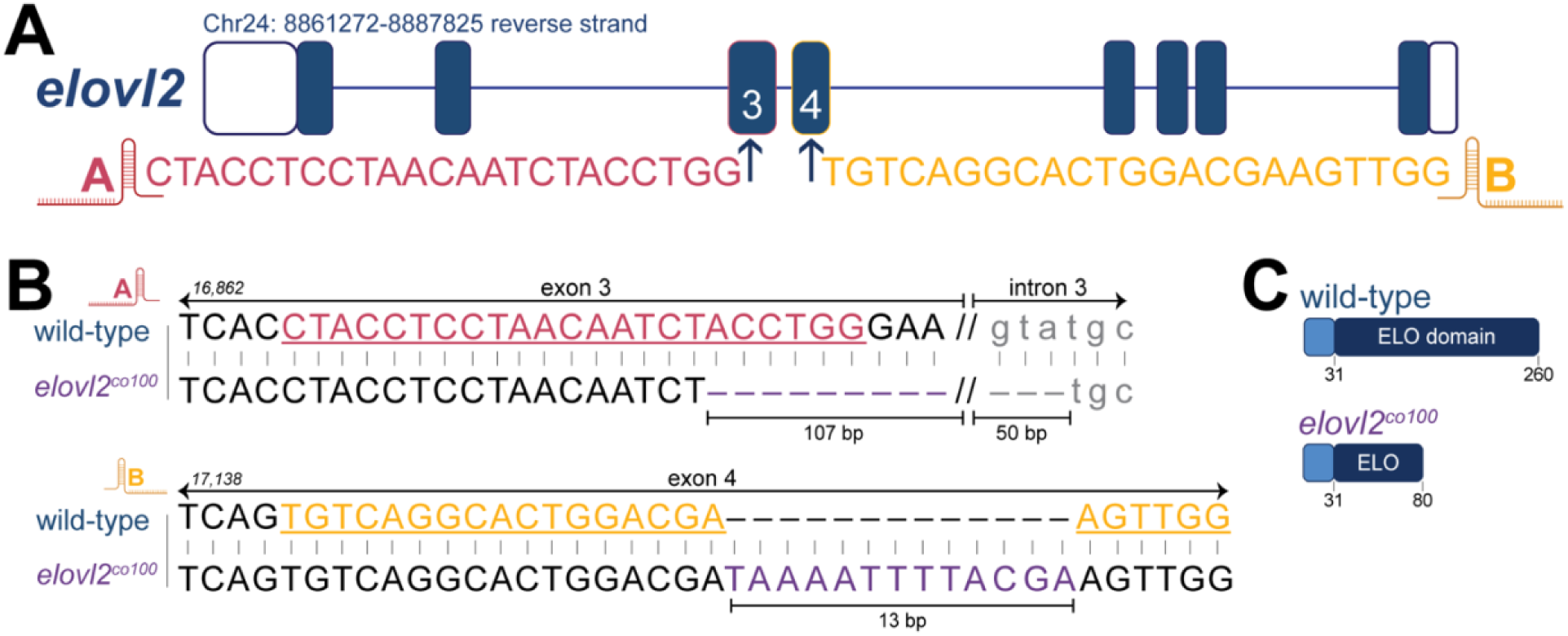
Generation of *elovl2* mutant zebrafish with CRISPR/Cas9 genome editing. **(A)** Schematic of the *elovl2* gene and gRNA sequences, identified as targets A and B. **(B)** Genomic sequences for wild-type *elovl2* and one of the two generated mutant alleles, *elovl2^co100^*. Sequence peak tracings and PCR verification are shown in **Supplemental Figure 2**. The *elovl2^co100^* allele has a 157 bp deletion (107 bp in exon 3; 50 bp in intron 3) and 13 bp insertion in exon 4. **(C)** Schematic of predicted Elovl2 protein in wild-type (260 AA) and *elovl2^co100^* (80 AA) fish. Genomic and protein results for the second generated mutant allele (*elovl2^co101^*) are shown in **Supplemental Figure 1**. All experiments were conducted using *elovl2^co100^* mutants unless otherwise noted.

### Maternal dietary DHA restriction and biosynthesis disruption depletes offspring DHA and increases ARA

Dietary DHA could override the need for endogenous DHA production via *elovl2*. To eliminate this possibility, we fed heterozygous *elovl2* mutant parents the DHA-deficient diet (DEF). We predicted that combining dietary DHA restriction and *elovl2* disruption at the maternal level would yield offspring with very low DHA status and thus provide an ideal model system.

To confirm that our maternal dietary and genetic manipulations altered offspring fatty acid profiles, we conducted targeted lipidomics analyses using whole embryos immediately after spawning (0 hpf) and at 4 dpf. Total DHA concentrations were significantly decreased in offspring produced by DEF-fed mothers compared to the SUFF group at both 0 hpf and 4 dpf (**Figure 5A**, **Tables 2 and 3**). Specifically, altering the maternal diet alone decreased 4 dpf WT offspring DHA levels by 41% (*P*<0.0001). This was surprising given that freshwater fish likely have a higher DHA biosynthesis capacity than humans (54), and that nutrient-restricted mothers often preferentially shuttle nutrients to their offspring at their own expense. DHA status was further decreased in 4 dpf *elovl2-*DEF offspring compared to WT-SUFF offspring (59% reduction, *P*<0.0001). *elovl2^+/+^* and *elovl2^-/-^* clutchmate siblings had similar DHA levels (*P=*0.716), suggesting that offspring DHA status largely depends on maternal factors. Like DHA, concentrations of the omega-3 LC-PUFA known as EPA were decreased in all DEF offspring groups at both developmental timepoints (**Tables 2 and 3**), which reflected maternal dietary EPA levels (**Table 1**).

**Figure 5.**
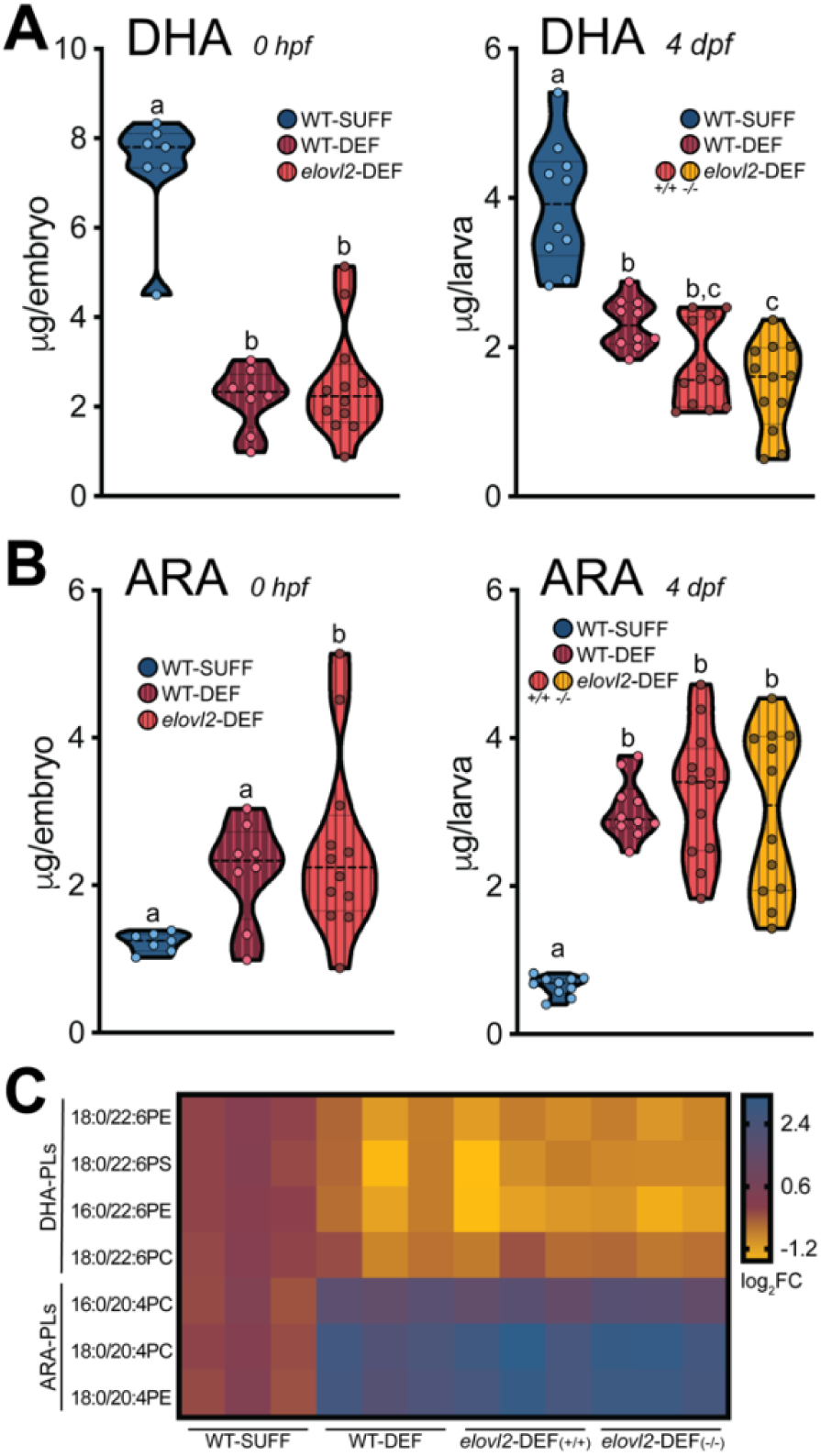
Maternal dietary DHA restriction and biosynthesis disruption depletes offspring DHA and increases ARA. Adult wild-type and *elovl2^+/-^*parents were fed either SUFF or DEF for 25-29 weeks prior to spawning. DHA **(A)** and ARA **(B)** concentrations in 0 hpf embryos and 4 dpf larvae measured by GC/MS. Offspring from *elovl2-*DEF mothers had similar fatty acid status, regardless of +/+ or -/- genotype. Dots represent individually analyzed offspring (n=6-12 fish/group). Groups labeled with different letters denote significant differences by 1-way ANOVA with Tukey’s post-hoc test (*P*<0.05). **(C)** Representative DHA-PL and ARA-PL species in 4 dpf larvae measured by LC/MS/MS (n=3 pooled replicates/group; n=3 larvae per pool). Values are log_2_(fold change) relative to the WT-SUFF group mean. DHA-PLs decreased and ARA-PLs increased in offspring from DEF-fed mothers compared to the WT-SUFF group. ARA, arachidonic acid; DHA, docosahexaenoic acid; PC, phosphatidylcholine; PE, phosphatidylethanolamine; PL, phospholipid; PS, phosphatidylserine.

**Table 2.** Embryo fatty acid profiles at 0 hpf (µg/embryo)*^a^*.

| fatty acid | WT-SUFF | WT-DEF | *elovl2*-DEF |
| --- | --- | --- | --- |
| 14:0 | 0.13 ± 0.02<sup>a</sup> | 0.06 ± 0.01<sup>b</sup> | 0.19 ± 0.12<sup>b</sup> |
| 16:0 | 7.3 ± 0.33 | 5.99 ± 0.54 | 6.93 ± 0.82 |
| 18:0 | 1.85 ± 0.06 | 2.02 ± 0.17 | 2.81 ± 0.34 |
| 16:1 | 0.11 ± 0.02<sup>a</sup> | 0.03 ± 0.01<sup>b</sup> | 0.05 ± 0.01<sup>b</sup> |
| 18:1 | 3.02 ± 0.27 | 2.91 ± 0.38 | 3.54 ± 0.46 |
| 18:2n-6 (LA) | 2.71 ± 0.35 | 4.39 ± 0.60 | 3.61 ± 0.48 |
| 18:3n-3 (ALA) | 0.24 ± 0.04 | 0.31 ± 0.04 | 0.22 ± 0.03 |
| 20:4n-6 (ARA) | 1.23 ± 0.05<sup>a</sup> | 3.46 ± 0.43<sup>a</sup> | 6.36 ± 1.08<sup>b</sup> |
| 20:5n-3 (EPA) | 2.63 ± 0.12<sup>a</sup> | 0.20 ± 0.03<sup>b</sup> | 0.17 ± 0.03<sup>b</sup> |
| 22:6n-3 (DHA) | 7.33 ± 0.49<sup>a</sup> | 2.18 ± 0.25<sup>b</sup> | 2.50 ± 0.36<sup>b</sup> |
| LA:ALA | 11.32 | 14.25 | 16.10 |
| ARA:DHA | 0.17 | 1.59 | 2.54 |

<sup>a</sup>fatty acids were analyzed by GC/MS using individual embryos (n=7-12 embryos/group). Values reported as mean ± SEM. For each fatty acid, different superscript letters denote significant differences by 1-way ANOVA with Tukey's post-hoc test, or when appropriate, Kruskal-Wallis test with Dunn's multiple comparisons test ( $P < 0.05$ ). Fatty acids without superscripts did not significantly differ between groups. ALA, alpha-linolenic acid; ARA, arachidonic acid; DEF, DHA-deficient diet; DHA, docosahexaenoic acid; EPA, eicosapentaenoic acid; LA, linoleic acid; SUFF, DHA-sufficient diet.

**Table 3.** Larval fatty acid profiles at 4 dpf (µg/larva)*^a^*.

| fatty acid | WT-SUFF | WT-DEF | *elovl2*-DEF (+/+) | *elovl2*-DEF (-/-) |
| --- | --- | --- | --- | --- |
| 14:0 | 0.07 ± 0.01<sup>a</sup> | 0.05 ± 0.01<sup>ab</sup> | 0.07 ± 0.01<sup>ab</sup> | 0.05 ± 0.01<sup>b</sup> |
| 16:0 | 3.52 ± 0.17 | 4.12 ± 0.19 | 3.63 ± 0.20 | 3.40 ± 0.30 |
| 18:0 | 1.26 ± 0.08<sup>a</sup> | 1.76 ± 0.11<sup>b</sup> | 2.04 ± 0.13<sup>b</sup> | 1.75 ± 0.13<sup>b</sup> |
| 16:1 | 0.04 ± 0.01<sup>a</sup> | 0.02 ± 0.01<sup>b</sup> | 0.02 ± 0.01<sup>b</sup> | 0.02 ± 0.01<sup>b</sup> |
| 18:1 | 1.01 ± 0.10 | 1.21 ± 0.09 | 1.23 ± 0.14 | 1.14 ± 0.14 |
| 18:2n-6 (LA) | 0.66 ± 0.09<sup>a</sup> | 1.35 ± 0.10<sup>b</sup> | 1.10 ± 0.13<sup>ab</sup> | 1.09 ± 0.15<sup>ab</sup> |
| 18:3n-3 (ALA) | 0.05 ± 0.01 | 0.07 ± 0.01 | 0.05 ± 0.01 | 0.05 ± 0.01 |
| 20:4n-6 (ARA) | 0.65 ± 0.04<sup>a</sup> | 3.04 ± 0.13<sup>b</sup> | 3.25 ± 0.26<sup>b</sup> | 2.99 ± 0.32<sup>b</sup> |
| 20:5n-3 (EPA) | 1.01 ± 0.07<sup>a</sup> | 0.13 ± 0.02<sup>b</sup> | 0.10 ± 0.01<sup>b</sup> | 0.09 ± 0.01<sup>b</sup> |
| 22:6n-3 (DHA) | 3.92 ± 0.26<sup>a</sup> | 2.32 ± 0.11<sup>b</sup> | 1.74 ± 0.16<sup>bc</sup> | 1.48 ± 0.17<sup>c</sup> |
| LA:ALA | 14.09 | 18.45 | 20.47 | 19.90 |
| ARA:DHA | 0.17 | 1.31 | 1.87 | 2.02 |

<sup>a</sup>fatty acids were analyzed by GC/MS using individual larvae (n=10-12 larvae/group). Values reported as mean ± SEM. For each fatty acid, different superscript letters denote significant differences by 1-way ANOVA with Tukey's post-hoc test, or when appropriate, Kruskal-Wallis test with Dunn's multiple comparisons test ( $P < 0.05$ ). Fatty acids without superscripts did not significantly differ by group. ALA, alpha-linolenic acid; ARA, arachidonic acid; DEF, DHA-deficient diet; DHA, docosahexaenoic acid; EPA, eicosapentaenoic acid; LA, linoleic acid; SUFF, DHA-sufficient diet.

Although there is minimal pre-formed DHA in DEF, this diet does contain its precursor, ALA (**Table 1**). Therefore, we also measured offspring ALA levels. Interestingly, ALA concentrations did not differ in embryos (**Table 2**) or larvae (**Table 3**) from DEF-fed mothers, despite approximately 2-fold higher dietary ALA levels in DEF compared to SUFF.

In contrast to the omega-3 PUFAs, two major omega-6 PUFAs (ARA and LA) were increased in offspring from DEF-fed mothers compared to WT-SUFF (**Figure 5B**, **Tables 2 and 3**). At 4 dpf, ARA levels were increased by 366% (*P*<0.0001) and 379% (*P*<0.0001) in WT-DEF and *elovl2-*DEF larvae, respectively. The high offspring ARA status in these groups is intriguing, given that the DEF maternal diet was deficient in ARA (**Table 1**). Overall, these lipidomics results imply a shift toward omega-6 LC-PUFA synthesis in the DEF maternal diet condition at the expense of omega-3 LC-PUFA synthesis, thus contributing to an omega-3 to omega-6 imbalance in offspring.

Although concentrations of saturated and unsaturated fatty acids, such as palmitic acid (16:0) and palmitoleic acid (16:1), differed in DEF compared to SUFF (**Table 1**), these fatty acids were relatively stable in offspring regardless of maternal diet. These results support our strategy to create a novel zebrafish model, as we have altered offspring omega-3 and omega-6 PUFAs without also substantially changing the profiles of other fatty acids.

Measuring fatty acid concentrations at both 0 hpf and 4 dpf also provides insight into how PUFA status changes during development. DHA levels, as well as most other analyzed fatty acids, were lower at 4 dpf compared to 0 hpf. This decrease with age is consistent with yolk lipids being used to meet the metabolic demands of the developing embryo.

DHA and ARA accumulate in the phospholipids (PL) of cell membranes, particularly in the eye and brain, where they contribute to the membrane’s physical properties. Notably, these PUFAs can also be released from the membrane to participate in signal transduction, both directly or indirectly through their bioactive derivatives (55). To confirm that DHA and ARA levels were altered in this CNS-relevant lipid class, we measured DHA-PL and ARA-PL species in pooled larvae at 4 dpf. The most abundant DHA-PL and ARA-PL species are shown in **Figure 4C**. In aggregate,

DHA-PLs were decreased by 44% (*P=*0.058) and 55% (*P*=0.004) in WT-DEF and *elovl2-*DEF larvae, respectively, compared to WT-SUFF. In contrast, the ARA-PLs were increased by 304% (*P*<0.0001) and 358% (*P*<0.0001) in WT-DEF and *elovl2-*DEF larvae, respectively. *elovl2^+/+^ and elovl2^-/-^* larvae in the *elovl2-*DEF group had similar levels of DHA-PLs (*P=*0.0.732) and ARA-PLs (*P=*0.897). Raw values for the DHA-PL and ARA-PL species used in these comparisons can be found in **Supplemental Table S5**. The offspring PL profiles align with our total fatty acid results. Altogether, our lipidomics findings demonstrate that we can generate zebrafish offspring with low DHA status by altering the maternal diet and disrupting *elovl2* gene function.

### DHA-restricted mothers produce offspring with a series of morphological phenotypes, and phenotype severity is related to offspring DHA and ARA status

While generating offspring for experiments, we observed a series of morphological phenotypes in WT-DEF and *elovl2-*DEF clutches (**Figure 6**). These clutches were produced by mothers fed DEF for 20-42 weeks. Four abnormal features were used to characterize the morphological phenotype and its severity. These features included: small eyes, curved axis, pericardial edema, and uninflated swim bladder. We binned larvae into the “severe” phenotype category if they exhibited ≥3 abnormal features. Some offspring exhibited only 1 to 3 of these features, but because we did not anticipate this morphological phenotype or its ranging severity before the study began, we were unable to systematically capture the number of offspring with all possible combinations of <3 features.

**Figure 6.**
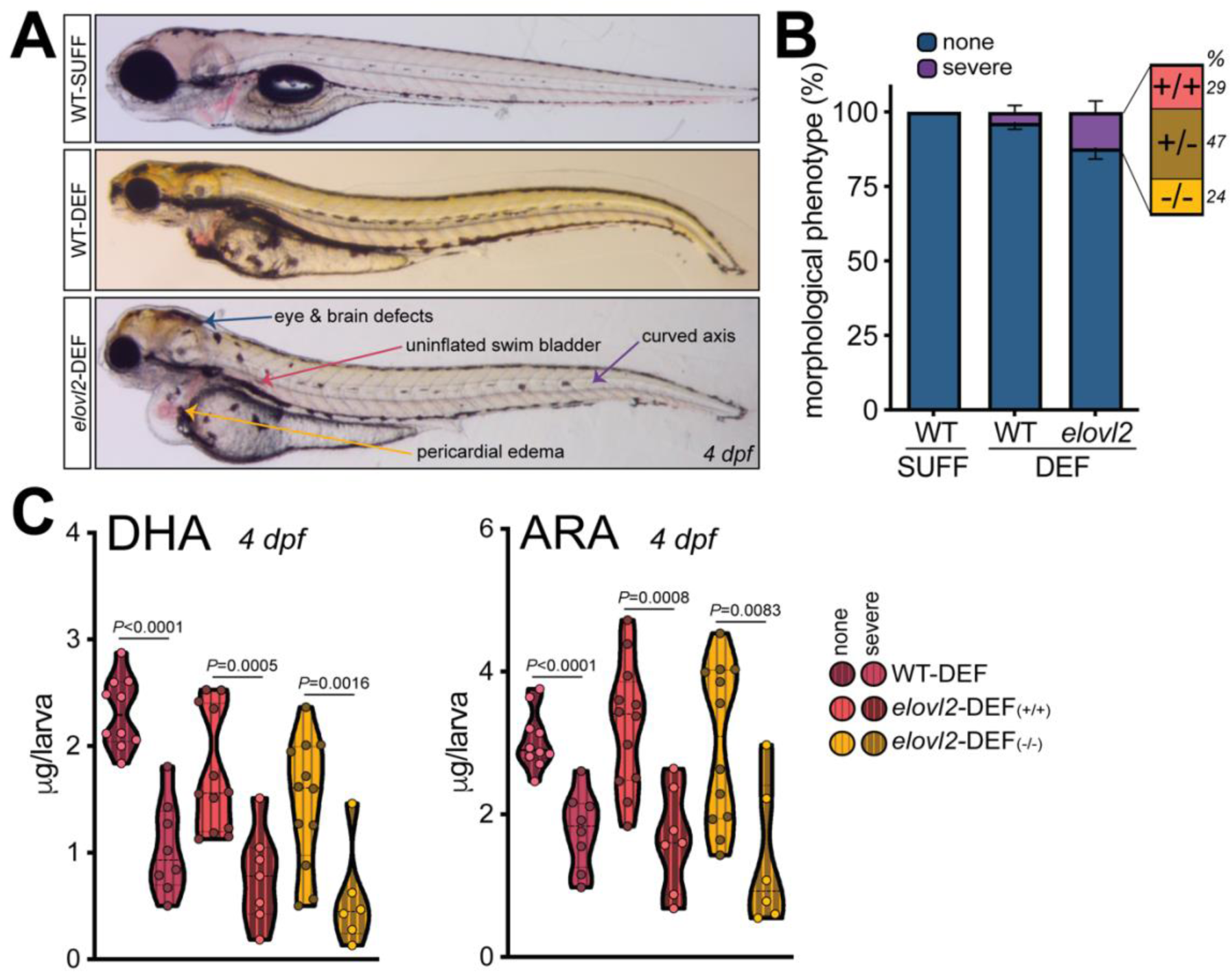
DHA-restricted mothers produce offspring with a series of morphological phenotypes, and phenotype severity is related to offspring DHA and ARA status. **(A)** Example 4 dpf larvae from each experimental group. The *elovl2*-DEF larva shown here is representative of the more severe morphological phenotype observed. This severe phenotype was characterized by the 4 abnormal features labeled in this panel. For both DEF groups, a series of phenotypes was observed between and within clutches, with some offspring exhibiting 1 to 3 of these features. For example, the WT-DEF larva in this panel has 3 of the 4 features (no pericardial edema). **(B)** Proportion of offspring per clutch with or without severe morphological abnormalities at 4 dpf, expressed as a mean percentage ± SEM (n=9-13 clutches/group, with n=998-1638 larvae evaluated/group). Larvae were binned into the “severe” phenotype category if they exhibited ≥3 of the following abnormal features: small eyes, curved axis, uninflated swim bladder, and pericardial edema. The severe phenotype was most often observed in *elovl2*-DEF offspring (12% per clutch), followed by WT-DEF (4%) and WT-SUFF (0%). The *elovl2-*DEF “severe” defects bar (purple) is magnified to show that offspring genotype does not explain the phenotype; the proportions of *elovl2* +/+ (29%) +/- (47%) and -/- (24%) larvae with severe defects follow the predicted 1:2:1 Mendelian ratios. Percentages were calculated from n=130 genotyped larvae collected at 4 dpf, generated from 7 *elovl2-*DEF clutches. **(C)** Offspring with severe phenotypes (≥3 abnormal features) have lower DHA and ARA status than their morphologically normal siblings. Dots represent individually analyzed larvae, generated from DEF-fed mothers for 20-42 weeks (n=6-12 larvae/group). Two-tailed unpaired Student’s *t-*tests or Mann-Whitney tests were used when appropriate.

The severe phenotype was most often observed in *elovl2*-DEF offspring (average of 12% per clutch), followed by WT-DEF (4%) and WT-SUFF (0%) (**Figure 6B**). Uninflated swim bladder was the only abnormal feature observed in WT-SUFF offspring (2% frequency). This is not unexpected because the swim bladder is still developing until ∼5 dpf. However, because uninflated swim bladders were observed in a much larger proportion of WT-DEF (11%) and *elovl2-*DEF (45%) offspring, this feature was included in our morphological assessment. The morphological phenotype was not explained by *elovl2-*DEF offspring genotype, as the proportions of *elovl2* +/+ (29%) +/- (47%) and -/- (24%) larvae with severe defects aligned with the predicted 1:2:1 Mendelian ratios (**Fig 6B**).

We observed some abnormal features (e.g., small eyes) as early as ∼24 hpf. A few 4 dpf *elovl2-*DEF larvae also exhibited what appeared to be blood in the brain, possibly indicative of blood brain barrier defects. However, we did not test blood brain barrier integrity in these larvae. In general, we observed both substantial intra-and inter-clutch variability in terms of morphological phenotype presence and severity. We speculate that this is due to differences in the composition of fatty acids deposited into each yolk by the mother.

To test whether the severe morphological phenotype was associated with altered omega-3 and omega-6 PUFA status, we conducted lipidomics analyses using representative WT-DEF and *elovl2*-DEF larvae at 4 dpf (**Figure 6C**). On average, DHA and ARA concentrations were decreased by ∼43% and ∼52%, respectively, compared to their morphologically normal siblings. Concentrations of other analyzed fatty acids, such as ALA, LA, and myristic acid (14:0), were changed to varying degrees in the abnormal larvae (**Supplemental Table S6**). Although not conclusive, these data support a link between the morphological phenotype and an imbalance in omega-3 and omega-6 LC-PUFA status in zebrafish offspring.

### Offspring with low DHA status have dysregulated vision and stress response gene pathways

To assess whole-body transcriptomic changes in our low DHA zebrafish model, we conducted bulk RNA-sequencing using pooled WT-SUFF and morphologically normal *elovl2-*DEF larvae at 4 dpf (**Figure 7**). Compared to WT-SUFF, 180 and 110 genes were down-and up-regulated, respectively, in the *elovl2-*DEF group (**Figure 7B**). Among the downregulated genes in the *elovl2-*DEF group was *elovl2* (-1.56 fold-change, q=1.8e-2), suggesting transcript clearance by nonsense mediated decay.

**Figure 7.**
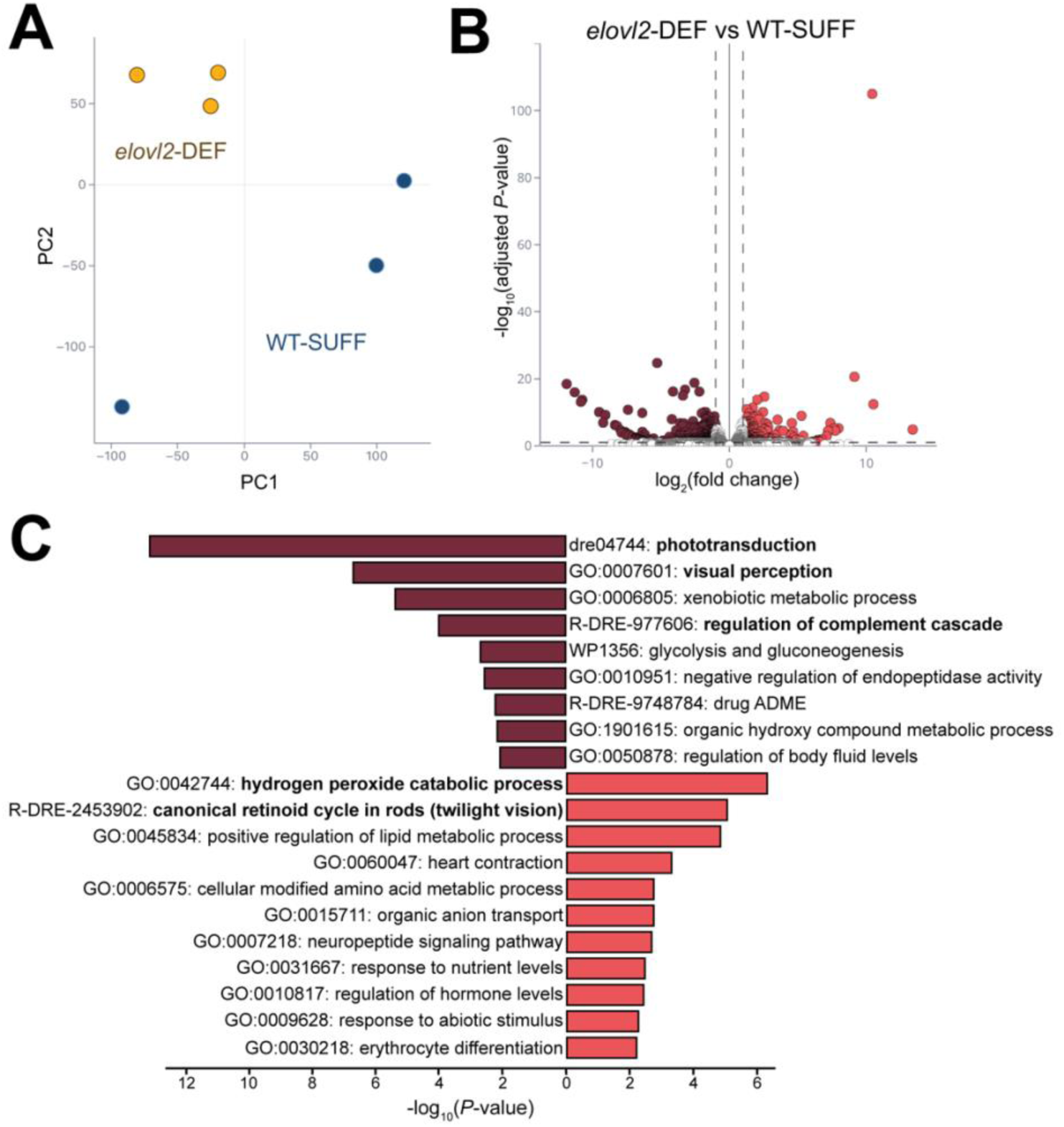
Offspring with low DHA status have dysregulated vision and stress response gene pathways. Morphologically normal 4 dpf larvae from wild-type mothers fed DHA-sufficient diet (WT-SUFF) or *elovl2^+/-^*mothers fed DHA-deficient diet (*elovl2-*DEF) were used for bulk RNA-seq (n=3 replicates/group, n=25 pooled whole larvae/replicate). **(A)** Principal component analysis showing that samples in the same group cluster together. **(B)** Volcano plot of differentially expressed genes in *elovl2-*DEF compared to WT-SUFF (290 genes). Dashed lines mark thresholds used to assess statistical significance (FDR-adjusted *P-*value *<*0.1, ≥ 2 fold change). **(C)** Significantly down-(maroon) and up-(coral) regulated gene pathways identified from a Metascape pathway enrichment analysis using the 290 DEGs (hypergeometric test with Benjamini-Hochberg procedure for multiple comparisons).

A pathway enrichment analysis revealed that the top dysregulated pathways were related to vision, stress response, and nutrient metabolism (**Figure 7C**). Given that we pooled whole larvae, the detection of differentially expressed genes within visual pathways is an exciting finding. Additionally, while a small eye morphological phenotype was observed in a proportion of their clutchmate siblings (**Figure 6**), the *elovl2-*DEF larvae selected for this experiment were morphologically normal heterozygotes. Two of the differentially expressed vision genes were *rho* (-2.01 fold-change, q=3.9e-5) and *rpe65a* (2.56 fold-change, q=1.3e-6), which encode rhodopsin and retinoid isomerohydrolase, respectively. Both proteins are essential components of the visual cycle of the retina. Rhodopsin is the light receptor in rod photoreceptor cells, and retinoid isomerohydrolase is critical for the regeneration of the visual pigment required for rod and cone-mediated vision (56).

The complement cascade is a key part of the innate immune response, as it facilitates the elimination of pathogens and damaged cells (57). Several genes involved in the regulation of complement cascade (*ela3l, c8g, and serping1*) were significantly downregulated in *elovl2-*DEF offspring. One of these genes, *serping1* (-27.29 fold-change, q=3.2e-6), encodes C1 inhibitor protein, which inhibits the initiation of the complement cascade (58). Downregulation of *serping1* may thereby lead to increased pathway activation and a heightened immune response.

The gene encoding cathepsin L, like (*ctsll*) was the highest upregulated gene (1951.68 fold-change, q=3.9e-128). Zebrafish cathepsin L, like is predicted to be involved in proteolysis as a cysteine-type endopeptidase localized to the lysosome. Interestingly, while our data are not CNS-specific, increased cysteine cathepsins have been implicated in the M1 polarization of microglia and neuroinflammation (59,60). Our RNA-seq results raise the possibility that low DHA status could increase offspring sensitivity to environmental insults and/or alter the immune response and microglia functions during neurodevelopment.

## Discussion

We created a novel larval zebrafish model with low DHA status by restricting maternal DHA intake and disrupting endogenous DHA production. Previous studies in rodents employed similar dietary and genetic approaches and demonstrated that low DHA status alters the immune response, synaptic plasticity markers, and cognitive performance (13,14,53,61). These findings underscore DHA’s critical role in brain development. However, we lack a deeper understanding of the cellular and molecular mechanisms involved, partially due to limitations of existing animal models. Addressing this fundamental knowledge gap is crucial for establishing DHA intake recommendations that support optimal brain development. Zebrafish offer a distinct advantage over rodents when investigating the cellular and molecular mechanisms guiding neurodevelopment. The zebrafish CNS develops rapidly post-fertilization, and larvae are transparent, enabling real-time visualization of individual CNS cells. Zebrafish are also increasingly used to study lipids, as the metabolic pathways and key proteins are well-conserved across vertebrates (62). Our innovative DHA-deficient zebrafish model will facilitate unprecedented explorations into DHA’s role in early life, unlocking new and exciting avenues in nutrition research.

To develop our model, we targeted Elovl2, an elongase enzyme in the DHA biosynthesis pathway that modulates systemic DHA levels in both mammals (18,19,41) and zebrafish (20,21,63). DHA metabolism primarily occurs in the liver, and thus hepatocytes highly express *elovl2* (42,64), but zebrafish, mouse, and human single-cell RNA-seq datasets also indicate that CNS astrocytes express *elovl2* (35,36,42–48). We confirmed *elovl2* expression profiles in larval brain and spinal cord astrocytes via RNA-FISH at 4 dpf. We also detected *elovl2* transcripts in the developing larval eye, consistent with a previous report in 3 dpf larvae (63). Our findings suggest that CNS cells have the enzymatic machinery required for local DHA synthesis. To further study local DHA production and utilization during neurodevelopment, an astrocyte-specific *elovl2* mutant could be a valuable tool. In the current study, we generated global *elovl2* mutants with the goal of depleting DHA in the whole embryo.

Select PUFAs, including DHA and ARA, accumulate in vertebrate reproductive tissues and are essential for both male and female fertility (65–69). Spermatogenesis is completely arrested in male *Elovl2^-/-^* mice (68), and female *elovl2^-/-^* zebrafish have impaired oocyte maturation and fecundity (69), demonstrating the need for endogenous PUFA synthesis in reproduction. To avoid these potential fertility issues, we incrossed *elovl2^+/-^* males and females to generate study offspring. Spawning success, clutch sizes, and embryo viability rates were similar between our *elovl2* mutant and wild-type experimental groups.

We showed that feeding DEF to wild-type mothers decreased offspring DHA levels by 41% compared to their SUFF counterparts, with further reductions (59%) in offspring from DEF-fed *elovl2^+/-^* mothers. Notably, the degree of DHA depletion in our *elovl2^-/-^* zebrafish larvae was modest compared to the 90% decrease observed in perinatal *Elovl2^-/-^* mice (18,19). Additionally, DHA levels were significantly lower in *Elovl2^-/-^* mice compared to their wild-type littermates, but in our study, levels were similar between *elovl2^-/-^* and *elovl2^+/+^* zebrafish siblings. Therefore, in mice, both the maternal and offspring *Elovl2* genotypes appear to contribute to DHA accretion (19), whereas maternal factors may dominate in zebrafish. We measured offspring fatty acid profiles after feeding DEF to mothers for ∼27 weeks, so it is also possible that more extreme DHA depletion could be achieved with longer maternal feeding periods. Offspring from DEF-fed mothers had a concomitant increase in ARA, a major omega-6 PUFA. Like omega-3 PUFAs, omega-6s are important constituents of cell membranes and regulate the immune response and inflammation (55). However, maintaining the proper balance between these lipid classes is critical because omega-3 and omega-6 PUFAs can elicit opposing effects; omega-3s are known for their anti-inflammatory properties, whereas omega-6s are considered pro-inflammatory in disease contexts (70). The optimal dietary omega-6 to omega-3 ratio for humans is debated but may range from 1:1 to 4:1 (71). A typical Western diet, in contrast, has a very high ratio of approximately 16:1, and this imbalance is associated with increased risk of cardiovascular disease, cancer, and immune conditions (70). The altered omega-6 to omega-3 ratio in our zebrafish offspring is therefore highly relevant to human health, and our model could be used to study multiple nutrition-associated chronic diseases.

Two factors related to our dietary and genetic manipulations may explain the increased offspring ARA levels. The same set of enzymes are used to bioconvert omega-6 and omega-3 precursors to the long-chain PUFAs, and this includes Elovl2. However, whereas Elovl2 elongates EPA (20:5n-3) to DPA (22:5n-3) prior to a desaturation step that yields DHA (22:6n-3), Elovl2 is not necessary for ARA production. Rather, Elovl2 elongates ARA (20:4n-6) to adrenic acid (22:4n-6). Therefore, the build-up of ARA could be a consequence of the *elovl2* mutation. Secondly, although the DEF diet does not contain pre-formed DHA or ARA, it does contain their respective precursors, ALA and LA. Due to the lipid source used in the DEF formulation (soybean oil), LA is approximately 2-fold higher than ALA in this diet. LA levels were also higher in offspring from DEF-fed mothers compared to SUFF-fed mothers. Therefore, a higher availability of LA in the maternal diet and embryo yolk likely prompted a shift toward ARA synthesis.

We measured fatty acid profiles in individual embryos rather than pooled offspring, which was a strength of our study. However, our lipidomics analyses captured whole-body rather than CNS-specific DHA levels, and thus we are assuming that CNS levels were also significantly lower. Our decreased offspring DHA-PL levels support this assumption since DHA-PLs are enriched in the brain and eye (72). Future studies should also consider measuring a panel of lipids derived from DHA and ARA that mediate inflammation, collectively known as oxylipins, to gain insight into the downstream consequences of altered PUFA status (73,74).

In contrast to rodents, a standardized reference diet for zebrafish is currently unavailable (38). However, efforts are underway to determine zebrafish nutrient requirements, including for DHA, which will eventually inform standardized diet formulations. Consequently, as our reference diet, we used a commercially available chow containing DHA, and we deemed this diet to be DHA-sufficient because it is known to support fish growth and reproduction (37). One limitation of our study is that we compared this reference chow diet to a nutrient-defined diet, and these diet types inherently differ in their ingredients as well as their fatty acid profiles beyond DHA. Nonetheless, this was a suitable dietary comparison for establishing our zebrafish model. Moving forward, mothers will be fed either the nutrient-defined DEF diet or a DHA-supplemented diet, in which high-purity DHA will replace up to 10% of the original lipid source (soybean oil). By transitioning mothers from DEF to a DHA-supplemented diet, we can also test whether maternal DHA supplementation rescues the offspring phenotype. Given that pregnant and lactating women are advised to supplement with DHA to support fetal and infant growth and development (75), these experiments have excellent translational potential.

We observed abnormal morphological phenotypes in offspring with low DHA status, characterized by small eyes, pericardial edema, curved body axis, and/or an uninflated swim bladder. Curiously, except for the small eyes, these abnormal features have also been reported in vitamin-E deficient zebrafish larvae (24,76), and both vitamin E-deficient (76) and *elovl2* crispant (63) zebrafish larvae exhibit impaired visual-motor responses when exposed to light/dark stimuli. The parallels between these larvae are intriguing because vitamin E is an antioxidant that protects brain PUFAs, including DHA, from lipid peroxidation (72), and vitamin E deficiency has been associated with long-chain PUFA depletion in zebrafish (23). In alignment with the visual behavior findings and other evidence that DHA prevents age-related vision loss (77), our bulk RNA-seq results provide support at the transcript level that DHA is critical for vision.

Pathways related to stress and immune response were also dysregulated in our RNA-seq dataset, possibly implicating microglia, the resident immune macrophages of the CNS. Research in perinatal mice suggests that low DHA exposure during development can disrupt the microglial immune response and increase microglial phagocytosis of synaptic elements (14). Interestingly, microglia also influence the formation and elimination of myelin, a major component of white matter that insulates axons and enhances nerve impulse conduction (15). Indeed, our lab’s recent zebrafish study revealed through *in vivo* time-lapse imaging that microglia can engulf excess myelin sheaths during normal development (16). Additionally, animals and preterm infants with low DHA status often exhibit white matter defects (78–80), but it is unknown how these features are connected. What cellular and molecular mechanisms could explain this link? Might low DHA status alter microglia functions, culminating in the abnormal regulation of myelination? Using our novel DHA-deficient zebrafish model, we can now start addressing fundamental neurodevelopmental questions. Importantly, these DHA investigations could significantly contribute to efforts that improve maternal and infant nutrition and prevent neurodevelopmental disorders.

## Abbreviations

ALA: alpha-linolenic acid
ARA: arachidonic acid
CNS: central nervous system
DEF: DHA-deficient diet
DHA: docosahexaenoic acid
*elovl2*: elongation of very long-chain fatty acid 2
EPA: eicosapentaenoic acid
RNA-FISH: RNA fluorescent *in situ* hybridization
LA: linolenic acid
PL: phospholipid
PUFA: polyunsaturated fatty acid
SUFF: DHA-sufficient diet

## Data availability statement

The data that support the findings of this study are available from the corresponding author upon reasonable request.

## Acknowledgements

We thank Rebecca O’Rourke for her assistance with the bulk RNA-seq analysis and the CU-AMC fish facility staff. We also thank the Colorado Nutrition Obesity Research Center (NORC) Lipidomics Core Laboratory (P30DK048520) and the University of Colorado Anschutz Medical Campus Cancer Center Genomics Shared Resource Core Facility (RRID:SCR_021984) (P30CA06934) for their contributions.

## CRediT author statement

*Katherine Ranard:* Conceptualization, Methodology, Formal analysis, Investigation, Writing-original draft, Writing-review & editing, Visualization, Funding acquisition. *Bruce Appel:* Conceptualization, Methodology, Writing-review & editing, Supervision, Funding acquisition.

## Funding sources

This work was supported by NIH/NINDS [F32NS131175 (KMR); R35NS122191 (B.A.)], NIH/NICHD [T32HD007186 (K.M.R)], and the Colorado Nutrition Obesity Research Center (NORC) Pilot & Feasibility Program [NIH/NIDDK, P30DK048520 (K.M.R)].

## Conflict of interest statement

The authors declare that they have no conflicts of interest with the contents of this article.

## Supplemental data

This article contains supplemental tables and figures.

## Notes

### Competing Interest Statement

The authors have declared no competing interest.

