## Supplemental materials for "Creation of a novel zebrafish model with low DHA status to study the role of maternal nutrition during neurodevelopment"

**Supplemental Table S1. DHA-sufficient diet (SUFF) composition^1^**

**ingredient**

fish meal

lecithin

wheat gluten

dried seaweed

fish oil

Maize starch

vitamin mix

mineral mix

**^1^**DHA-sufficient diet (SUFF) is Gemma Micro commercial lab diet. The ingredient list was obtained from Skretting.

**Supplemental Table S2. DHA-deficient diet (DEF) composition^1^**

| **ingredient** | **g/100 g** |
| --- | --- |
| wheat gluten | 15 |
| casein | 30.5 |
| egg whites | 4 |
| cellulose | 3 |
| vitamin mix^2^ | 4 |
| mineral mix^3^ | 4 |
| modified food starch (National Starch Food Innovation) | 26.5 |
| soybean oil | 7 |
| lecithin (Lipoid PC 18:0, Lipoid LLC) | 5 |
| Stay C (Argent Chemical Laboratories) | 1 |
| alpha-tocopherol acetate (Archer Daniels Midland) | 0.05 |

**^1^**Unless otherwise noted, ingredients were sourced from Dyets Inc (Bethlehem, PA). Diets were prepared at Oregon State University Sinnhuber Aquatic Research Laboratory. The DHA-deficient diet (DEF) was formulated as described in Lebold *et al,* 2011 (https://doi.org/10.3945/jn.111.144279), except the current study did not use tocopherol-stripped versions of soybean oil and lecithin.

**^2^**Vitamin mix contained the following (g/kg vitamin mix): vitamin A (500,000 IU/g), 0.15; vitamin D_3_(400,000 United States Pharmacopeia (USP)/mg), 6.2445; vitamin K, 0.025; thiamine, 0.15; riboflavin, 0.25; vitamin B_6_, 0.125; pantothenic acid, 0.75; niacin, 1.25; biotin, 0.005; folate, 0.05; vitamin B_12_, 0.0005; myoinositol, 6.25; para-amino benzoic acid (PABA), 1; celufil (α-cellulose), 983.75.

**^3^**Mineral mix contained the following (g/kg mineral mix): calcium carbonate, 19.23; calcium phosphate dibasic (2H_2_0), 766.29; citric acid, 5.28; cupric carbonate, 0.36; ferric citrate, 2.99; magnesium oxide, 22.89; manganese carbonate, 5.65; sodium chloride, 28.02; disodium hydrogen phosphate, 11.89; zinc carbonate, 0.97; potassium phosphate dibasic, 74.16; potassium sulfate, 62.26; potassium iodide, 0.01.

**Supplemental Table S3**. **CRISPR/Cas9 reagents and genotyping primers**

| **resource type** | **sequences & additional information** |
| --- | --- |
| cRNA (Dr.Cas9.ELOVL2.1.AA), IDT | 5’-CTACCTCCTAACAATCTACC-3’ (exon 3) |
| cRNA (Dr.Cas9.ELOVL2.1.AB), IDT | 5’-TGTCAGGCACTGGACGAAGT-3’ (exon 4) |
| target A genotyping primers | forward, 5'-GTTTGTTTGATGTCAGATACCCG-3';  reverse, 5'-CCAAAC ACATGAAGCACAGC-3' |
| target B genotyping primers | forward, 5'-CTGTGCTTCATGTGTTTGGTAG-3';  reverse, 5'-ACATGAGTGCAATCCAGATGT-3' |
| *elovl2^co100^* allele genotyping primers | target A:  forward, 5'-GTTTGTTTGATGTCAGATACCCG-3;  reverse, 5'-CCAAACACATGAAGCACAGC-3'  (amplicon sizes: wild-type, 314 bp; mutant, 157 bp);  target B:  forward, 5'-CTGTGCTTCATGTGTTTGGTAG-3';  reverse, 5'-AGCATCAGTATTTTCACGAGCA-3'  (amplicon sizes: wild-type, 176 bp; mutant, 189 bp) |
| *elovl2^co101^* allele genotyping primers | target A:  forward, 5'-TGCTGGATTCCTACACACCA-3’;  reverse, 5'-TGCTGGTCTGTTTCTCATGT-3'  (amplicon sizes: wild-type, 92 bp; mutant, 88 bp);  target B:  forward, 5'-CTGTGCTTCATGTGTTTGGTAG-3';  reverse, 5'-AGCATCAGTATTTTCACGAGCA-3'  (amplicon sizes: wild-type, 176 bp; mutant, 166 bp) |

**Supplemental Table S4. Fatty acid concentrations are similar between *elovl2^co100^* and *elovl2^co101^* offspring (µg/larva)^1^**

| **fatty acid** | **co100** | | **co101** | |
| --- | --- | --- | --- | --- |
|  | ***+/+*** | ***-/-*** | ***+/+*** | ***-/-*** |
| 14:0 | 0.07 ± 0.01 | 0.05 ± 0.01 | 0.06 ± 0.01 | 0.09 ± 0.03 |
| 16:0 | 3.63 ± 0.20 | 3.40 ± 0.30 | 3.73 ± 0.23 | 3.50 ± 0.35 |
| 18:0 | 2.04 ± 0.13^a^ | 1.75 ± 0.13^ab^ | 1.45 ± 0.09^b^ | 1.45 ± 0.19^b^ |
| 16:1 | 0.02 ± 0.00 | 0.02 ± 0.01 | 0.02 ± 0.01 | 0.02 ± 0.01 |
| 18:1 | 1.23 ± 0.14 | 1.14 ± 0.14 | 1.12 ± 0.11 | 0.94 ± 0.12 |
| 18:2n-6 (LA) | 1.10 ± 0.13 | 1.09 ± 0.15 | 1.09 ± 0.12 | 0.78 ± 0.16 |
| 18:3n-3 (ALA) | 0.05 ± 0.01 | 0.05 ± 0.01 | 0.06 ± 0.01 | 0.03 ± 0.01 |
| 20:4n-6 (ARA) | 3.25 ± 0.26 | 2.99 ± 0.32 | 2.67 ± 0.24 | 2.56 ± 0.33 |
| 20:5n-3 (EPA) | 0.10 ± 0.01 | 0.09 ± 0.01 | 0.09 ± 0.01 | 0.07 ± 0.01 |
| 22:6n-3 (DHA) | 1.74 ± 0.16 | 1.48 ± 0.17 | 1.55 ± 0.14 | 1.22 ± 0.14 |
| LA:ALA | 20.47 | 19.90 | 19.55 | 28.80 |
| ARA:DHA | 1.87 | 2.02 | 1.72 | 2.10 |

^1^fatty acids were analyzed by GC/MS using individual 4 dpf larvae generated from heterozygous *elovl2* parents fed DEF for 20-29 weeks prior to spawning (n=7-12 larvae/group). Values reported as mean ± SEM. For each fatty acid, a 2-way ANOVA test was used to assess significant differences for the main effects of allele (co100 vs. co101) and larval genotype (+/+ vs. -/-). There was a significant effect of allele for 18:0 (*P=*0.0019), and superscript letters denote differences identified from Sidak’s multiple comparisons test. There were no significant differences between groups for the other analyzed fatty acids. ALA, alpha-linolenic acid; ARA, arachidonic acid; DEF, DHA-deficient diet; DHA, docosahexaenoic acid; EPA, eicosapentaenoic acid; LA, linoleic acid.

**Supplemental Table S5**. **DHA-PL and ARA-PL profiles in WT-SUFF, WT-DEF, and *elovl2-*DEF offspring at 4 dpf (pmol/sample)^1^**

| **PL species** | **WT-SUFF** | **WT-DEF** | ***elovl2-*DEF (+/+)** | ***elovl2-*DEF (-/-)** |
| --- | --- | --- | --- | --- |
| **DHA-PLs** |  |  |  |  |
| **18:0/22:6PS** | 445.66 ± 34.00 | 251.77 ± 48.20 | 219.09 ± 35.12 | 246.72 ± 1.40 |
| **16:0/22:6PE** | 266.26 ± 10.06 | 155.36 ± 19.82 | 113.14 ± 11.08 | 119.97 ± 8.53 |
| **18:0/22:6PE** | 470.86 ± 23.39 | 284.83 ± 34.54 | 254.97 ± 20.73 | 262.56 ± 17.26 |
| **16:0/22:6PC** | 1313.19 ± 37.06 | 705.27 ± 53.89 | 475.80 ± 24.40 | 506.58 ± 26.42 |
| **18:0/22:6PC** | 222.67 ± 14.92 | 158.67 ± 20.95 | 161.52 ± 14.91 | 151.55 ± 5.43 |
| **18:1/22:6PC** | 153.26 ± 7.10 | 100.49 ± 12.34 | 84.57 ± 9.15 | 79.91 ± 8.04 |
| **ARA-PLs** |  |  |  |  |
| **18:0/20:4PS** | 11.01 ± 1.78 | 34.79 ± 6.01 | 40.87 ± 5.86 | 46.07 ± 1.62 |
| **16:0/20:4PE** | 26.90 ± 3.28 | 117.16 ± 14.63 | 110.03 ± 10.96 | 120.64 ± 8.22 |
| **18:0/20:4PE** | 72.02 ± 8.65 | 352.46 ± 41.74 | 455.57 ± 54.03 | 467.06 ± 27.65 |
| **16:0/20:4PC** | 259.46 ± 32.72 | 833.76 ± 55.30 | 787.17 ± 73.94 | 880.42 ± 79.83 |
| **18:0/20:4PC** | 56.51 ± 4.44 | 293.02 ± 29.17 | 398.46 ± 60.64 | 410.26 ± 38.59 |
| **18:1/20:4PC** | 33.11 ± 3.77 | 167.97 ± 15.86 | 190.22 ± 15.54 | 200.38 ± 12.55 |

^1^phospholipid species were analyzed by LC/MS/MS using pooled 4 dpf larvae (n=3 replicates/group, with n=3 larvae pooled/sample). Values are mean ± SEM. ARA, arachidonic acid; DEF, DHA-deficient diet; DHA, docosahexaenoic acid; EPA, eicosapentaenoic acid; PE, phosphatidylethanolamine; PC, phosphatidylcholine; PL, phospholipid; PS, phosphatidylserine; SUFF, DHA-sufficient diet.

**Supplemental Table S6**. **Fatty acid profiles in WT-DEF and *elovl2-*DEF offspring with severe morphological phenotypes (µg/larva)^1^**

| **fatty acid** | **WT-DEF** | ***elovl2-*DEF (+/+)** | ***elovl2-*DEF (-/-)** |
| --- | --- | --- | --- |
| 14:0 | 0.05 ± 0.01 | 0.33 ± 0.27 | 0.06 ± 0.01 |
| 16:0 | 2.80 ± 0.31 | 3.65 ± 0.98 | 2.12 ± 0.40 |
| 18:0 | 1.22 ± 0.15 | 1.62 ± 0.19 | 1.14 ± 0.17 |
| 16:1 | 0.04 ± 0.02 | 0.02 ± 0.01 | 0.01 ± 0.01 |
| 18:1 | 0.83 ± 0.11 | 0.95 ± 0.23 | 0.67 ± 0.23 |
| 18:2n-6 (LA) | 0.83 ± 0.14 | 0.96 ± 0.34 | 0.68 ± 0.24 |
| 18:3n-3 (ALA) | 0.04 ± 0.01 | 0.07 ± 0.03 | 0.04 ± 0.01 |
| 20:4n-6 (ARA) | 1.68 ± 0.20 | 1.77 ± 0.28 | 1.37 ± 0.41 |
| 20:5n-3 (EPA) | 0.05 ± 0.01 | 0.06 ± 0.02 | 0.04 ± 0.01 |
| 22:6n-3 (DHA) | 0.97 ± 0.15 | 0.83 ± 0.19 | 0.57 ± 0.19 |
| LA:ALA | 18.83 | 14.81 | 17.92 |
| ARA:DHA | 1.71 | 2.13 | 2.42 |

^1^fatty acids were analyzed by GC/MS individual larvae (n=10-12 larvae/group). Values reported as mean ± SEM.

ALA, alpha-linolenic acid; ARA, arachidonic acid; DEF, DHA-deficient diet; DHA, docosahexaenoic acid; EPA, eicosapentaenoic acid; LA, linoleic acid; SUFF, DHA-sufficient diet.

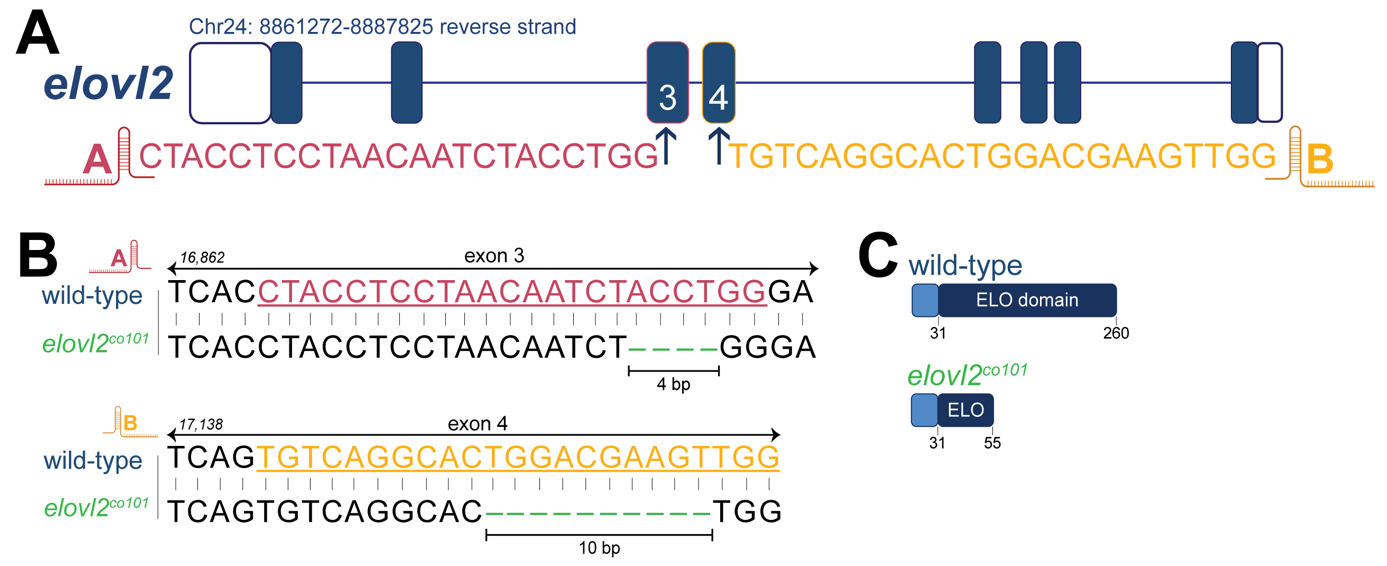

**Supplemental Figure S1. Generation of *elovl2^co101^* mutant zebrafish with CRISPR/Cas9 genome editing. (A)** Schematic of the *elovl2* gene and gRNA sequences, identified as targets A and B. **(B)** Genomic sequences for wild-type *elovl2* and the second generated mutant allele (*elovl2^co101^*), which has 4 and 10 bp deletions in exons 3 and 4, respectively. **(C)** Schematic of predicted Elovl2 protein in wild-type (260 AA) and *elovl2^co101^* (55 AA) fish.

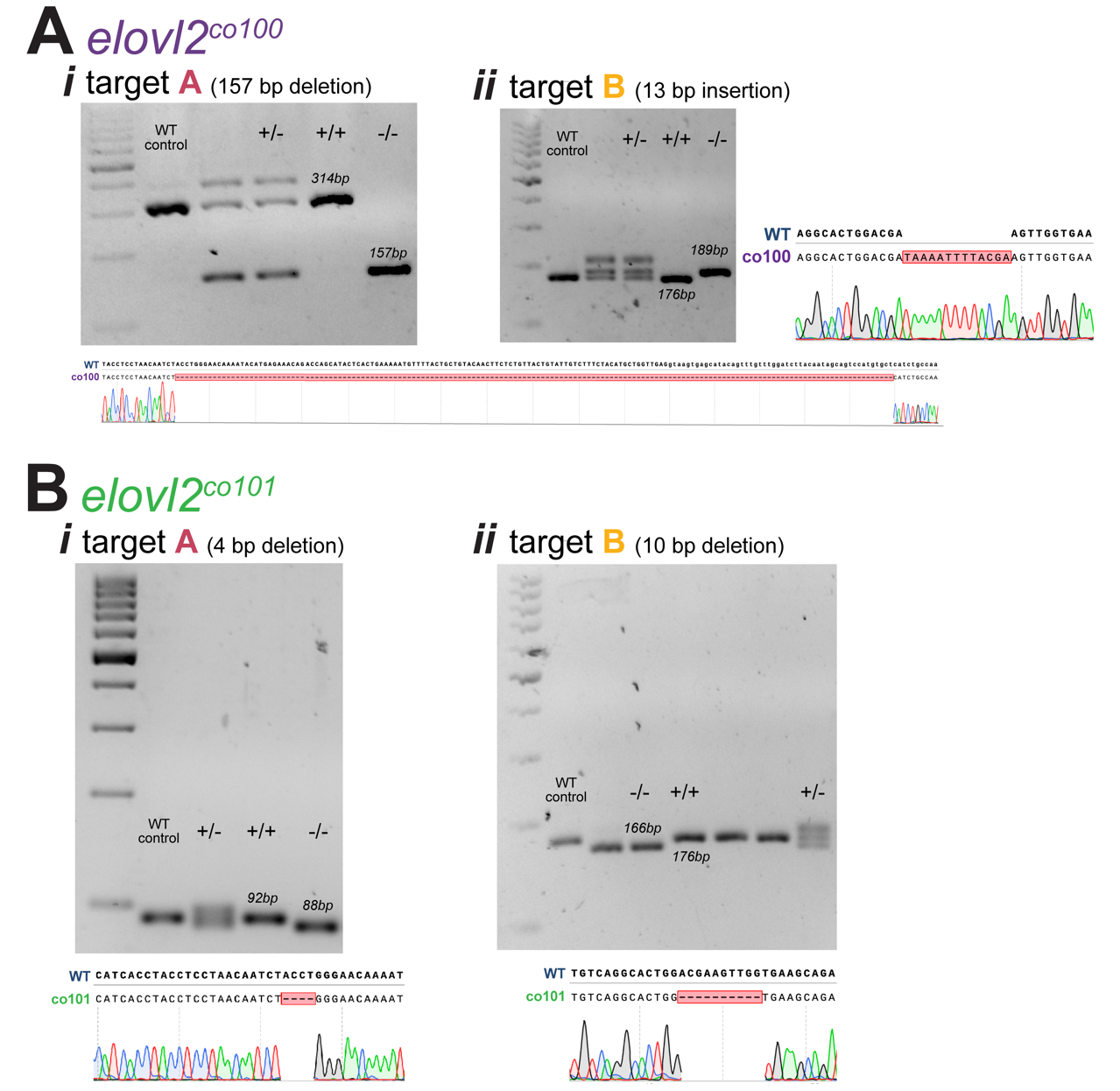

**Supplemental Figure S2. Genomic validation of (A) *elovl2^co100^* and (B) *elovl2^co101^* mutant alleles.** Genotyping results for PCR-amplified regions ran on a 3% agarose gel and Sanger sequencing peak tracings for each mutant allele’s (*i)* target A and (*ii)* target B sites. All gels show a GeneRuler 100bp Plus DNA ladder (lane 1) and a wild-type negative control sample (lane 2). The sequence alignments and mutant allele peak tracing diagrams were acquired from SnapGene.

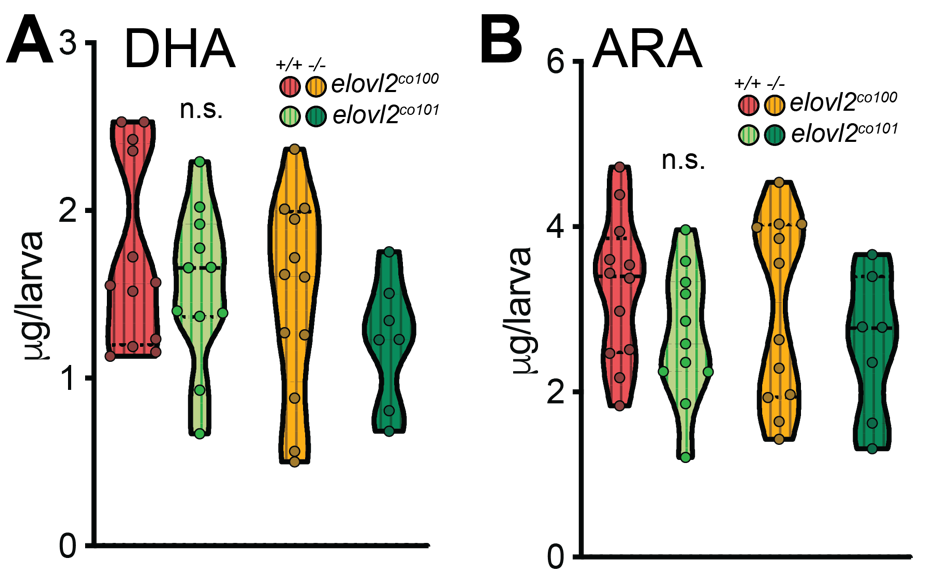

**Supplemental Figure S3. DHA and ARA levels are similar in *elovl2^co100^* and *elovl2^co101^* offspring.** Adult heterozygous *elovl2^co100^* and *elovl2^co101^* parents were fed DEF for 20-29 weeks prior to spawning. DHA **(A)** and ARA **(B)** concentrations in 4 dpf larvae measured by GC/MS. Larvae generated from the co100 and co101 mutant lines have similar fatty acid status. Dots represent individually analyzed offspring (n=7-12 fish/group). The main effects of allele (co100 vs. co101) and larval genotype (+/+ vs. -/-) did not differ by 2-way ANOVA. ARA, arachidonic acid; DHA, docosahexaenoic acid; n.s., not significant.
